# High lumenal chloride in the lysosome is critical for lysosome function

**DOI:** 10.1101/141788

**Authors:** Kasturi Chakraborty, KaHo Leung, Yamuna Krishnan

## Abstract

Lysosomes are organelles responsible for the breakdown and recycling of cellular machinery. Dysfunctional lysosomes give rise to lysosomal storage disorders as well as common neurodegenerative diseases. Here, we use a DNA-based, fluorescent chloride reporter to measure lysosomal chloride in *Caenorhabditis elegans* as well as murine and human cell culture models of lysosomal diseases. We find that the lysosome is highly enriched in chloride, and that chloride reduction correlates directly with a loss in the degradative function of the lysosome. In nematodes and mammalian cell culture models of diverse lysosomal disorders, where previously only lysosomal pH dysregulation has been described, massive reduction of lumenal chloride is observed that is ~10^4^-fold greater than the accompanying pH change. Reducing chloride within the lysosome impacts Ca^2+^ release from the lysosome and impedes the activity of specific lysosomal enzymes indicating a broader role for chloride in lysosomal function. −149 words.

## Introduction

Chloride is the most abundant, soluble anion in the body. Cytosolic chloride can be as low as ~ 45 mM, while extracellular chloride is ~110 mM (Treharne et al., 2006),(Sonawane et al., 2002). Chloride concentration values thus span a wide range and yet, in each compartment, it is quite tightly regulated (Sonawane and Verkman, 2003). For example, in early endosomes it is ~40 mM, late endosomes it is ~70 mM and lysosomes it is ~108 mM (Hara-Chikuma et al., 2005; Saha et al., 2015; Sonawane et al., 2002). Chloride levels are stringently regulated by chloride channels such as cystic fibrosis transmembrane regulator (CFTR), the CLC family of channels or calcium activated chloride channels, and their dysregulation is directly linked to several diseases including cystic fibrosis, myotonia, epilepsy, hyperekplexia or deafness (Planells-Cases and Jentsch, 2009). Chloride is largely considered to function as a counter ion only to balance changes in cation fluxes related to signaling (Scott and Gruenberg, 2011). In one form, this balancing function serves to reset the membrane potential of depolarized neurons through the operation of plasma membrane resident chloride channels/exchangers (Chen, 2005). In another form, it serves to continuously facilitate organelle acidification, through the operation of intracellular chloride channels (Stauber and Jentsch, 2013). Despite its importance in cell function, intracellular chloride has never been visualized or quantitated *in vivo*. Here, using a previously developed, pH-independent, DNA-based fluorescent chloride reporter called *Clensor*, we have made the first measure of chloride in a live multicellular organism, creating *in vivo* chloride maps of lysosomes in *C. elegans*.

Our investigations reveal that lysosomal chloride levels *in vivo* are even higher than extracellular chloride levels. Others and we have shown that lysosomes have the highest lumenal acidity and the highest lumenal chloride respectively, among all endocytic organelles (Saha et al., 2015; Weinert et al., 2010). Although lumenal acidity has been shown to be critical to the degradative function of the lysosome (Appelqvist et al., 2013; Eskelinen et al., 2003), the necessity for such high lysosomal chloride is unknown. In fact, in many lysosomal storage disorders, lumenal hypoacidification compromises the degradative function of the lysosome leading to the toxic build-up of cellular cargo targeted to the lysosome for removal, resulting in lethality (Guha et al., 2014). Lysosomal storage disorders (LSDs) are a diverse collection of ~70 different rare, genetic diseases that arise due to dysfunctional lysosomes (Samie and Xu, 2014). Dysfunction in turn arises from mutations that compromise protein transport into the lysosome, the function of lysosomal enzymes, or lysosomal membrane integrity (Futerman and van Meer, 2004). Importantly, for a sub-set of lysosomal disorders e.g., osteopetrosis or neuronal ceroid lipofuscinoses (NCL), lysosomal hypoacidification is not observed (Kasper et al., 2005). Both these conditions result from a loss of function of the lysosomal H^+^-Cl^−^ exchange transporter CLC-7 (Kasper et al., 2005). In both mice and flies, lysosomal pH is normal, yet both mice and flies were badly affected (Poët et al., 2006; Weinert et al., 2010).

The lysosome performs multiple functions due to its highly fusogenic nature. It fuses with the plasma membrane to bring about plasma membrane repair as well as lysosomal exocytosis, it fuses with the autophagosome to bring about autophagy, it is involved in nutrient sensing and it fuses with endocytic cargo to bring about cargo degradation (Appelqvist et al., 2013; Xu and Ren, 2015). To understand which, if any, of these functions is affected by chloride dysregulation, we chose to study genes related to osteopetrosis in the versatile genetic model organism *Caenorhabditis elegans*. By leveraging the DNA scaffold of *Clensor* as a natural substrate along with its ability to quantitate chloride, we could simultaneously probe the degradative capacity of the lysosome *in vivo* and then in cultured mammalian cells. Our findings reveal that depleting lysosomal chloride showed a direct correlation with loss of the degradative function of the lysosome. We found that lowering lysosomal chloride also reduced the level of Ca^2+^ released from the lysosome. We also observed that reduction of lysosomal chloride inhibited the activity of specific lysosomal enzymes such as cathepsin C and arylsulfatase B. The role of chloride in defective lysosomal degradation has been hypothesized in the past (Stauber and Jentsch, 2013; Wartosch and Stauber, 2010; Wartosch et al., 2009), and our studies provide the first mechanistic proof of a broader role for chloride in lysosome function.

## Results and Discussion

### Reporter design and uptake pathway in coelomocytes of *C. elegans*

In this study we use two DNA nanodevices, called the I-switch and *Clensor*, to fluorescently quantitate pH and chloride respectively (Modi et al., 2009; Saha et al., 2015). The I-switch is composed of two DNA oligonucleotides. One of these can form an i-motif, which is an unusual DNA structure formed by protonated cytosines (Gehring et al., 1993). In the I-switch intrastrand i-motif formation is used to bring about a pH-dependent conformational change, that leverages fluorescence resonance energy transfer (FRET) to create a ratiometric fluorescent pH reporter. (Figure 1; figure supplementary 2)

The DNA-based chloride sensor, *Clensor*, is composed of three modules: a sensing module, a normalizing module and a targeting module (Figure 1a) (Saha et al., 2015),(Prakash et al., 2016). The sensing module is a 12 base long peptide nucleic acid (PNA) oligomer conjugated to a fluorescent, chloride-sensitive molecule 10,10′-Bis[3-carboxypropyl]-9,9′-biacridinium dinitrate (BAC), (Figure 1a)(Sonawane et al., 2002). The normalizing module is a 38 nt DNA sequence bearing an Alexa 647 fluorophore that is insensitive to Cl^−^. The targeting module is a 26 nt double stranded DNA domain that targets it to the lysosome via the endolysosomal pathway by engaging the scavenger receptor or ALBR pathway. In physiological environments, BAC specifically undergoes collisional quenching by Cl^−^, thus lowering its fluorescence intensity (G) linearly with increasing Cl^−^ concentrations. In contrast, the fluorescence intensity of Alexa 647 (R) remains constant (Figure 1b). This results in R/G ratios of *Clensor* emission intensities varying linearly with [Cl^−^] over the entire physiological regime of [Cl^−^]. Since the response of *Clensor* is insensitive to pH changes, it enables the quantitation of lumenal chloride in organelles of living cells regardless of their lumenal pH(Saha et al., 2015).

**Figure 1:**
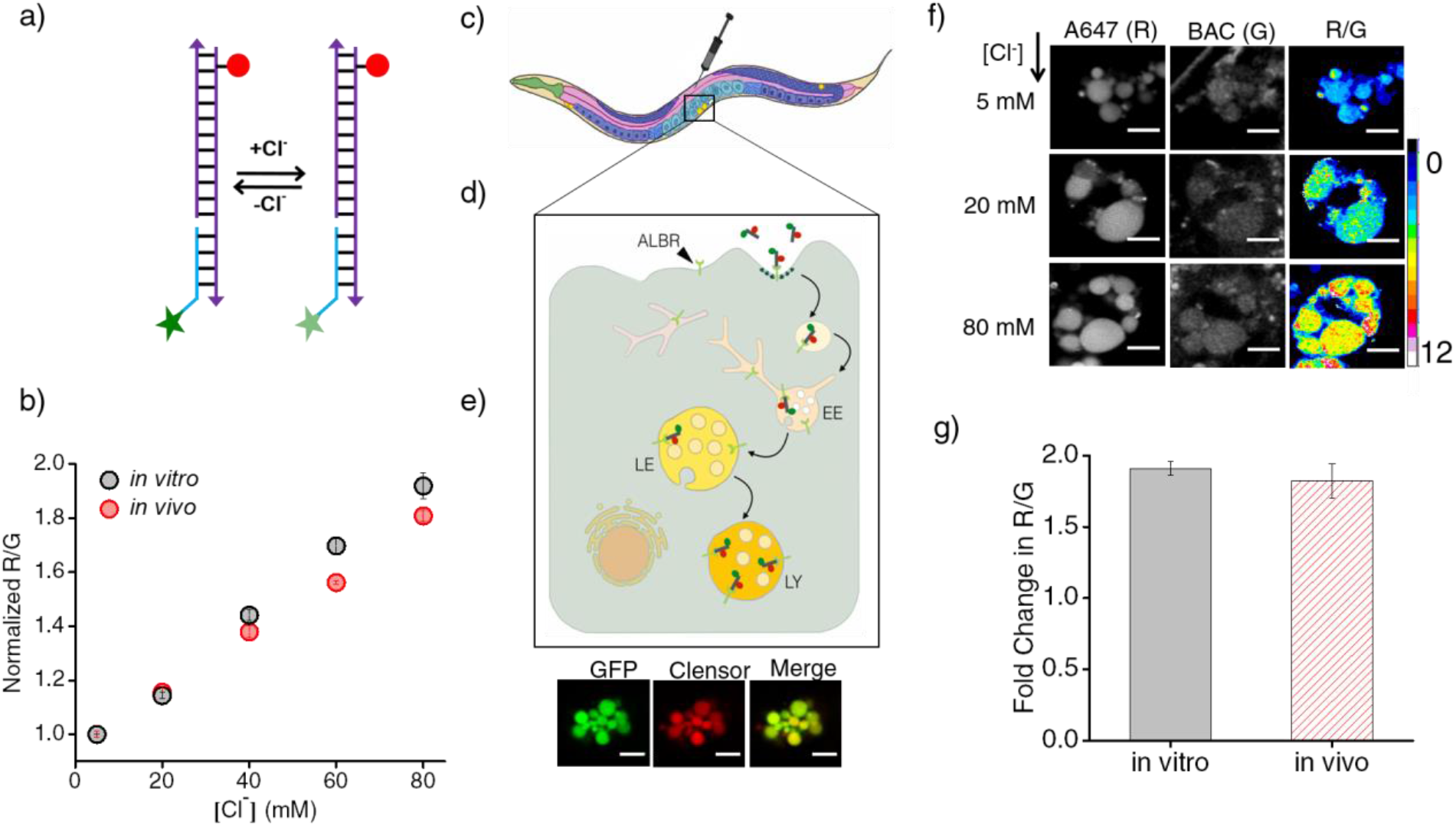
*Clensor* recapitulates its chloride sensing characteristics *in vivo*. (a) Schematic of the DNA nanodevice *Clensor*, which functions as a ratiometric fluorescent reporter for the chloride ion (Cl^−^). It bears a Cl^−^ sensitive fluorophore, BAC (green star) and a Cl^−^ insensitive fluorophore, Alexa 647 (red circle) (b) Calibration profile of *Clensor in vitro* (red) and *in vivo* (black) given by normalized Alexa 647 (R) and BAC (G) intensity ratios versus [Cl^−^]. (c) Receptor mediated endocytic uptake of *Clensor* in coelomocytes post injection in *C. elegans*. (d) *Clensor* is trafficked by the anionic ligand binding receptor (ALBR) from the early endosome (EE) to the late endosome (LE) and then lysosome (LY). (e) Image showing colocalization of *Clensor_A647_* (red) microinjected in the pseudocoelom with coelomocytes marked by GFP (green). Scale bar: 5 μm. (f) Representative images of endosomes in coelomocytes labeled with *Clensor* and clamped at the indicated chloride concentrations ([Cl^−^]). Images are acquired in the Alexa 647 (R) and BAC (G) channels from which corresponding R/G images are shown in pseudocolor. The *in vivo* calibration profile is shown in (b). Scale bar: 5 μm. Error bars indicate s.e.m. (n = 15 cells, ≥50 endosomes) (g) *In vitro* and *in vivo* fold change in R/G ratios of *Clensor* from 5 mM to 80 mM [Cl^−^].

### Targeting *Clensor* to lysosomes of coelomocytes in *C. elegans*

Coelomocytes of *C. elegans* are known to endocytose foreign substances injected in the body cavity(Fares and Greenwald, 2001). The polyanionic phosphate backbone of DNA can be co-opted to target it to scavenger receptors and thereby label organelles on the endolysosomal pathway in tissue macrophages and coelomocytes in *C. elegans* (Figure 1c and d) (Modi et al., 2009; Saha et al., 2015; Surana et al., 2011). Alexa 647 labelled I-switch (I4^cLY^) and *Clensor* were each injected in the pseudocoelom of 1-day-old adult worms expressing *pmyo-3*::ssGFP. In these worms, soluble GFP synthesized in muscles and secreted into the pseudocoelom is actively internalized by the coelomocytes resulting in GFP labeling of the coelomocytes(Fares and Greenwald, 2001). After 1 h, both devices quantitatively colocalize with GFP indicating that they specifically mark endosomes in coelomocytes (Figure 1e and Figure 1; figure supplementary 1c). Endocytic uptake of DNA nanodevices was performed in the presence of 30 equivalents of maleylated bovine serum albumin (mBSA), a well-known competitor for the anionic ligand binding receptor (ALBR) pathway (Gough and Gordon, 2000). Coelomocyte labeling by I4^cLY^or *Clensor* were both efficiently competed out by mBSA indicating that both reporters were internalized by ALBRs and trafficked along the endolysosomal pathway (Figure 1; figure supplementary 1b)(Surana et al., 2011).

### *In vivo* performance of DNA reporters

Next, the functionality of I4^cLY^ and *Clensor* were assessed *in vivo*. To generate an *in vivo* calibration curve for the I-switch I4^cLY^, coelomocytes labeled with I4^cLY^ were clamped at various pH values between pH 4 and 7.5 as described previously and in the supporting information(Surana et al., 2011). This indicated that, as expected, the I-switch showed *in vitro* and *in vivo* performance characteristics that were extremely well matched (Figure 1; figure supplementary 2 b-e). To assess the *in vivo* functionality of *Clensor*, a standard Cl^−^ calibration profile was generated by clamping the lumenal [Cl^−^] to that of an externally added buffer containing known [Cl^−^] as described previously for cultured cells (Saha et al., 2015). Endosomes of coelomocytes were labeled with *Clensor* and fluorescence images were acquired in the BAC channel (G) as well as Alexa 647 channel (R) as described (see methods), from which were obtained R/G ratios of every endosome clamped at a specific [Cl^−^] (Figure 1b). Endosomal R/G ratios showed a linear dependence on [Cl^−^] with ~2-fold change in R/G values from 5 mM to 80 mM [Cl^−^] (Figure 1f and g). This is very well matched with its *in vitro* fold change in R/G over the same regime of [Cl^−^].

### DNA nanodevices localize specifically in lysosomes in diverse genetic backgrounds

Before performing quantitative chloride imaging in various mutant nematodes, we checked whether lysosomal targeting of *Clensor* and the I-switch were preserved in a variety of genetic backgrounds of our interest. *Clensor* was injected into LMP-1::GFP worms treated with RNAi against specific lysosomal storage disorder (LSD)-related genes or genes linked to osteopetrosis. We observed significant colocalization (>74%) of *Clensor* with LMP-1-GFP labeled lysosomes in these coelomocytes (Figure 3; figure supplementary 2 b and c). Given that both *Clensor* and the I-switch robustly labeled lysosomes of coelomocytes in wild type worms (N2), mutants and RNAi knockdowns of a range of LSD-related genes, we explored whether these devices could report on alterations, if any, in the lumenal ionicity in these lysosomes, and thereby possibly report on lysosome dysfunction.

### Quantitative *in vivo* imaging of chloride in lysosomes in *C. elegans*

As an initial study, we focused on *C. elegans* nematodes in which genes related to osteopetrosis are mutated. Osteopetrosis results from non-functional osteoclasts that lead to increased bone mass and density due to a failure in bone resorption (Sobacchi et al., 2013). In humans, osteopetrosis results from mutations in a lysosomal chloride-proton antiporter CLCN7, and its auxiliary factor OSTM1 (Figure 2a) (Kornak et al., 2001; Lange et al., 2006). It also results from mutations in TCIRG1, which is the a3 subunit of a lysosomal V-ATPase (Kornak et al., 2000) and SNX-10, a sorting nexin implicated in lysosome transport to form the ruffled border of osteoclasts, which is critical for osteoclast function (Aker et al., 2012)(Figure 2a). Lysosomes of CLC7 knockout mice show normal lumenal pH, yet the mice manifest osteopetrosis as well as neurodegeneration, indicating that despite the apparently normal lumenal milieu, the organelle is still dysfunctional (Kasper et al., 2005). The *C. elegans* homologs for these genes are *clh-6* (CLCN7), *F42A8.3 (ostm-1;* OSTM1), *unc-32* (TCIRG1) and *snx-3* (SNX10) (Figure 2a).

**Figure 2:**
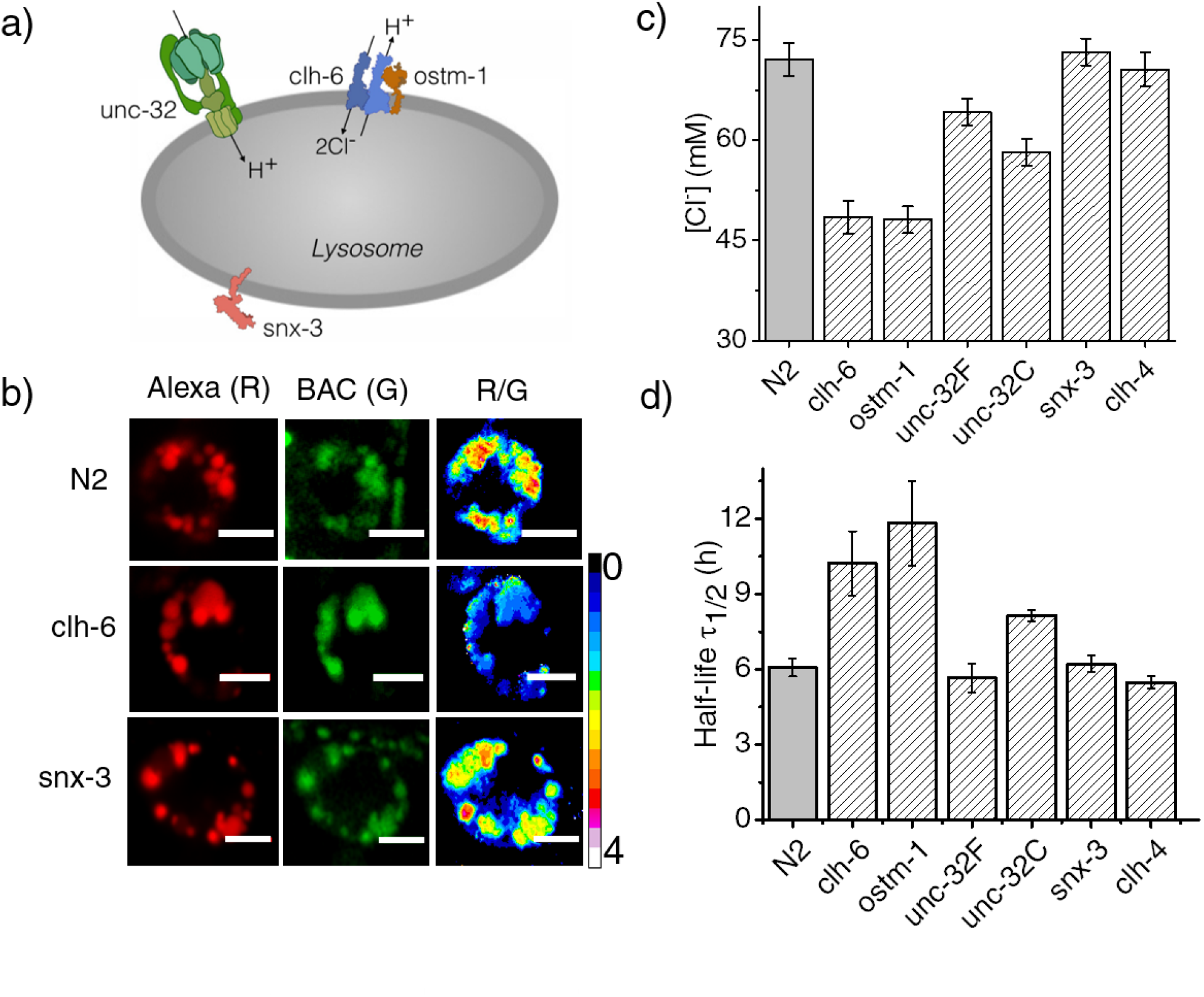
Dysregulation in lysosomal [Cl^−^] correlates with reduced lysosomal degradation. (a) Schematic depicting protein players involved in autosomal recessive osteopetrosis. (b) Representative images of *Clensor* in lysosomes of coelomocytes, in the indicated genetic backgrounds acquired in the Alexa 647 (R) and BAC (G) channels and their corresponding pseudocolored R/G images. Scale bar, 5 μm. (b) Lysosomal chloride concentrations ([Cl^−^]) as measured using *Clensor* in indicated genetic background (n = 10 worms, ≥100 lysosomes). (d) Degradative capacity of lysosomes of coelomocytes of nematodes with the indicated genetic backgrounds as given by the measured half-life of *Clensor*. Error bars indicate s.e.m.

*Clensor* was injected into N2, *clh-6* and *unc-32* mutants and RNAi knockdowns of *ostm-1* and *snx-3*. Chloride concentrations in the lysosomes of each genetic background at 60 min post injection were obtained (Figure 2 b,c and Figure 2; figure supplementary 1). N2 worms showed a chloride concentration of ~75 mM. Knocking down *clh-6* and *ostm-1* resulted in a dramatic decrease of lysosomal chloride to ~45 mM due to the loss of function in chloride transport. Lumenal pH in the lysosomes of these mutants was normal, consistent with findings in both flies and mice(Saha et al., 2015; Weinert et al., 2010). As a control, knocking down a plasma membrane resident CLC channel such as *clh-4* showed no effect on either lysosomal chloride or pH (Schriever et al., 1999). *unc-32c* is a non-functional mutant of the V-ATPase *a* sub-unit, while *unc-32f* is a hypomorph(Pujol et al., 2001). Interestingly, a clear inverse correlation with *unc-32* functionality was obtained when comparing their lysosomal chloride levels i.e., ~55 mM and ~65 mM for *unc-32c* and *unc-32f* respectively. Importantly, *snx-3* knockdowns showed lysosomal chloride levels that mirrored those of wild type lysosomes. In all genetic backgrounds, we observed that lysosomal chloride concentrations showed no correlation with lysosome morphology (Figure 3; figure supplementary 1d).

### Reducing lumenal chloride lowers the degradative capacity of the lysosome

Dead and necrotic bone cells release their endogenous chromatin extracellularly - thus duplex DNA constitutes cellular debris and is physiologically relevant cargo for degradation in the lysosome of phagocytic cells (Elmore, 2007; Luo and Loison, 2008). Coelomocytes are phagocytic cells of *C. elegans*, and thus, the half-life of *Clensor* or I4^cLY^ in these cells constitutes a direct measure of the degradative capacity of the lysosome(Tahseen, 2009). We used a previously established assay to measure the half-life of I-switches in lysosomes(Surana et al., 2013). Worms were injected with 500 nM I4^cLY^ and the fluorescence intensity obtained in 10 cells at each indicated time point was quantitated as a function of time. The I-switch I4^cLY^ had a half-life of ~6 h in normal lysosomes, which nearly doubled when either *clh-6* or *ostm-1* were knocked down (Figure 2d and Figure 2; figure supplementary 2). Both *unc-32c* and *unc-32f* mutants showed near-normal lysosome degradation capacity, inversely correlated with their lysosomal chloride values (Figure 2d and Figure 2; figure supplementary 2).

In this context, data from *snx-3* and *unc-32f* mutants support that high lysosomal chloride is critical to the degradation function of the lysosome. In humans, SNX10 is thought to be responsible for the vesicular sorting of V-ATPase from the Golgi or for its targeting to the ruffled border(Aker et al., 2012). Non-functional SNX10 can thus be considered a ‘secondary V-ATPase deficiency’, phenocopying a V-ATPase deficiency and showing osteoclasts without ruffled borders due to defective lysosomal transport (Aker et al., 2012). Importantly, lysosomal pH in *snx-3* knockdowns was compromised by 0.3 pH units, while that in *unc-32* knockdowns was compromised by 0.2 pH units (Figure 2 figure supplementary 1) (Chen et al., 2012). Yet both these genetic backgrounds showed completely normal lysosomal degradation capacity, that is consistent with their normal lumenal chloride levels, rather than their defective pH levels. This further supports that high lysosomal chloride is a sensitive correlate of the degradative function of the lysosome.

### Lysosomal chloride is highly depleted in lysosomal storage disorders

Since lysosomal chloride dysregulation correlated with a loss of degradative ability of the lysosome, we wondered whether the converse was true, i.e., whether lysosomes known to be defective in degradation as seen in lysosomal storage disorders, showed depleted chloride levels. Given that in higher organisms such as mice and humans, high acidity has also been shown to be essential for proper lysosome function (Mindell, 2012), we measured both lysosomal pH and lysosomal chloride in *C. elegans* mutants and RNAi knockdowns for a range of genes that are known to cause lysosomal storage disorders. These included a selection of diseases due to dysfunctional enzymes that metabolize sugar derivatives, such as mannose and glycosaminoglycans, as well as lipids such as sphingomyelin and glucosylceramide. Lysosomal pH and chloride measurements were made with I4^cLY^ and *Clensor* respectively, in each genetic background at 60 min post injection (Figure 3a and b). We found that in C. *elegans* mutants for Gaucher’s disease, Batten disease, different forms of NCL, MPS VI and Niemann Pick A/B disease, lysosomal chloride levels were severely compromised (Figure 3a and b). Dysfunctional lysosomes showed three types of ion profiles, those where either lysosomal acidity or chloride levels were reduced, and those where both lysosomal acidity and chloride were reduced. The magnitude of proton dysregulation in these defective lysosomes ranged between 1.9 – 2.8 μM. However, the magnitude of lysosomal chloride showed a stark drop, decreasing by 19 – 34 mM in most mutants. Importantly, in mammalian cell culture models for many of these diseases e.g., Gaucher’s disease, NCL, MPS VI, etc., only pH dysregulation has been reported (Bach et al., 1999; Holopainen et al., 2001; Sillence, 2013). Yet we find that in *C. elegans* models of these diseases that chloride levels are highly compromised. Chloride decreases by nearly four orders of magnitude more than proton decrease, and the percentage changes of both ions are similar.

**Figure 3:**
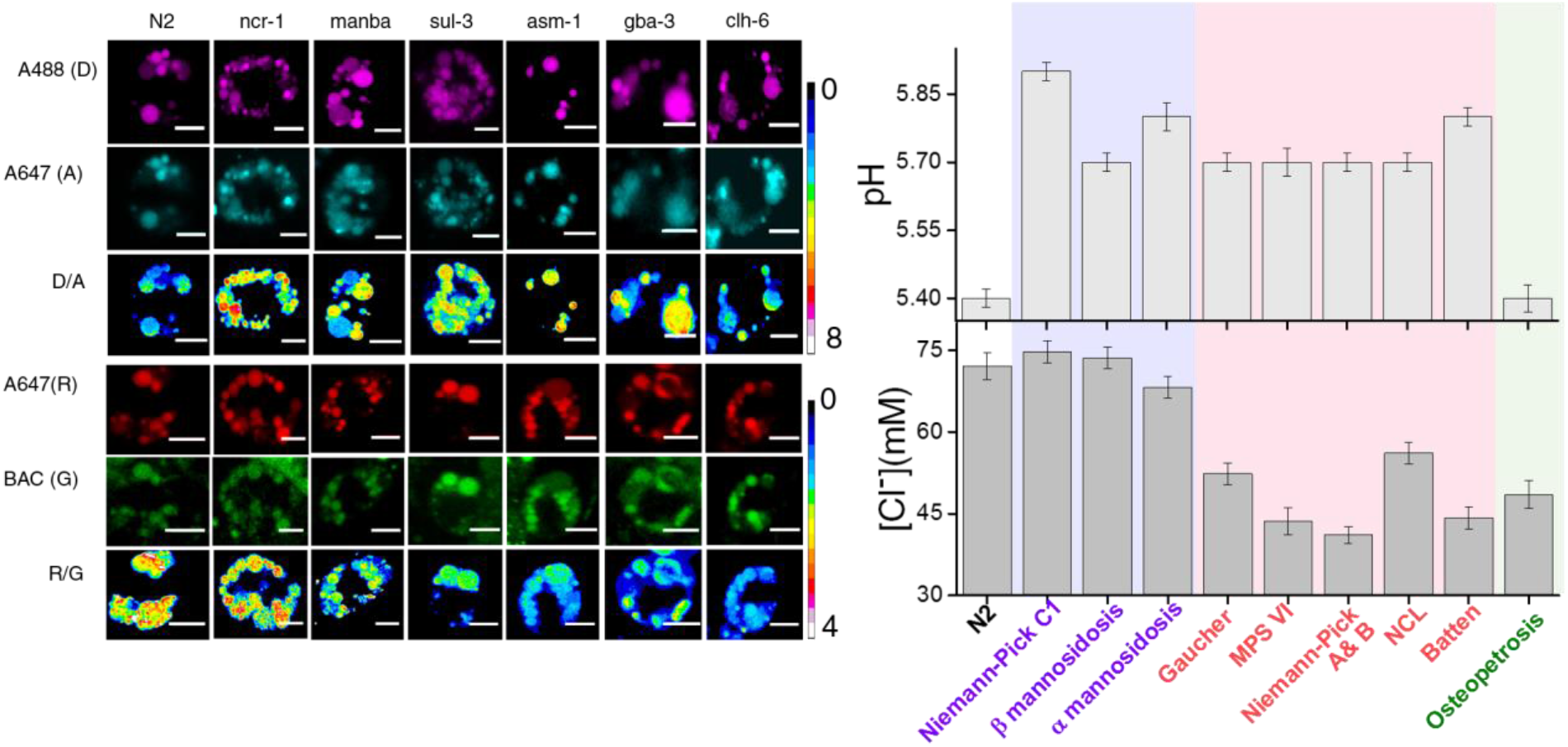
Lysosomal chloride dysregulation is observed in nematode models in several pH-related lysosomal storage disorders. Representative pH maps of lysosomes in coelomocytes labelled with (a) a DNA-based pH reporter, I4^cLY^_A488/A647_, in the indicated genetic backgrounds. Images were acquired in the donor (D, magenta) and acceptor (A, cyan) channels and the corresponding pseudocolored D/A images. (b) Representative [Cl^−^] maps of lysosomes acquired in these genetic backgrounds using *Clensor*. Images are acquired in the Alexa 647 (R) and BAC (G) channels and the corresponding pseudocolored R/G images are shown. Scale bar, 5 μm. (c) Quantification of lysosomal pH and lysosomal chloride in *C. elegans* mutants or RNAi knockdowns of genes responsible for the indicated lysosomal storage diseases in humans. Mutants are grouped according to dysregulation only in lysosomal pH (purple box); only in lysosomal chloride (green box) and both lysosomal pH and chloride (pink box) for n = 10 worms (≥100 lysosomes) Error bars indicate s.e.m.

To check whether such chloride decrease is observed also in higher organisms, we made pH and chloride measurements in mammalian cell culture models of two relatively common lysosomal storage disorders. Macrophages are a convenient cell culture system to study lysosomal storage disorders as they can be isolated from blood samples and have a lifetime of 3 weeks in culture(Vincent et al., 1992). We re-created two widely used murine and human cell culture models of Gaucher’s disease by inhibiting β-glucosidase with its well-known inhibitor conduritol β epoxide (CBE) in murine and human macrophages namely, J774A.1 and THP-1 cells respectively (Hein et al., 2013, 2007; Schueler et al., 2004). We also recreated common mammalian cell culture models of Niemann-Pick A/B disease by inhibiting acid sphinogomyelinase (SMPD1) in J774A.1 and THP-1 cells with a widely used inhibitor amitriptyline hydrochloride (AH)(Aldo et al., 2013; Jones et al., 2008). First we confirmed that *Clensor* and our DNA-based pH reporter localized exclusively in lysosomes. In both cell lines, DNA nanodevices (500 nM) were uptaken from the extracellular milieu by the scavenger receptors, followed the endolysosomal pathway and showed quantitative colocalization with lysosomes that were pre-labelled with TMR-Dextran (Figure 4 figure supplementary 3 a and b). In-cell calibration curves of both pH (Figure 4 figure supplementary 1) and chloride reporters (Figure 4a) were well matched with their *in vitro* calibration profiles, indicating that both sensor integrity and performance were quantitatively preserved at the time of making lysosomal pH and chloride measurements in these cells. Both human and murine lysosomes in normal macrophages showed chloride concentrations close to ~118 mM, revealing that lysosomes have the highest chloride levels compared to any other endocytic organelle (Saha et al., 2015; Sonawane et al., 2002). This is nearly 10-15% higher than even extracellular chloride concentrations, which reaches only up to 105-110 mM (Arosio and Ratto, 2014).

**Figure 4:**
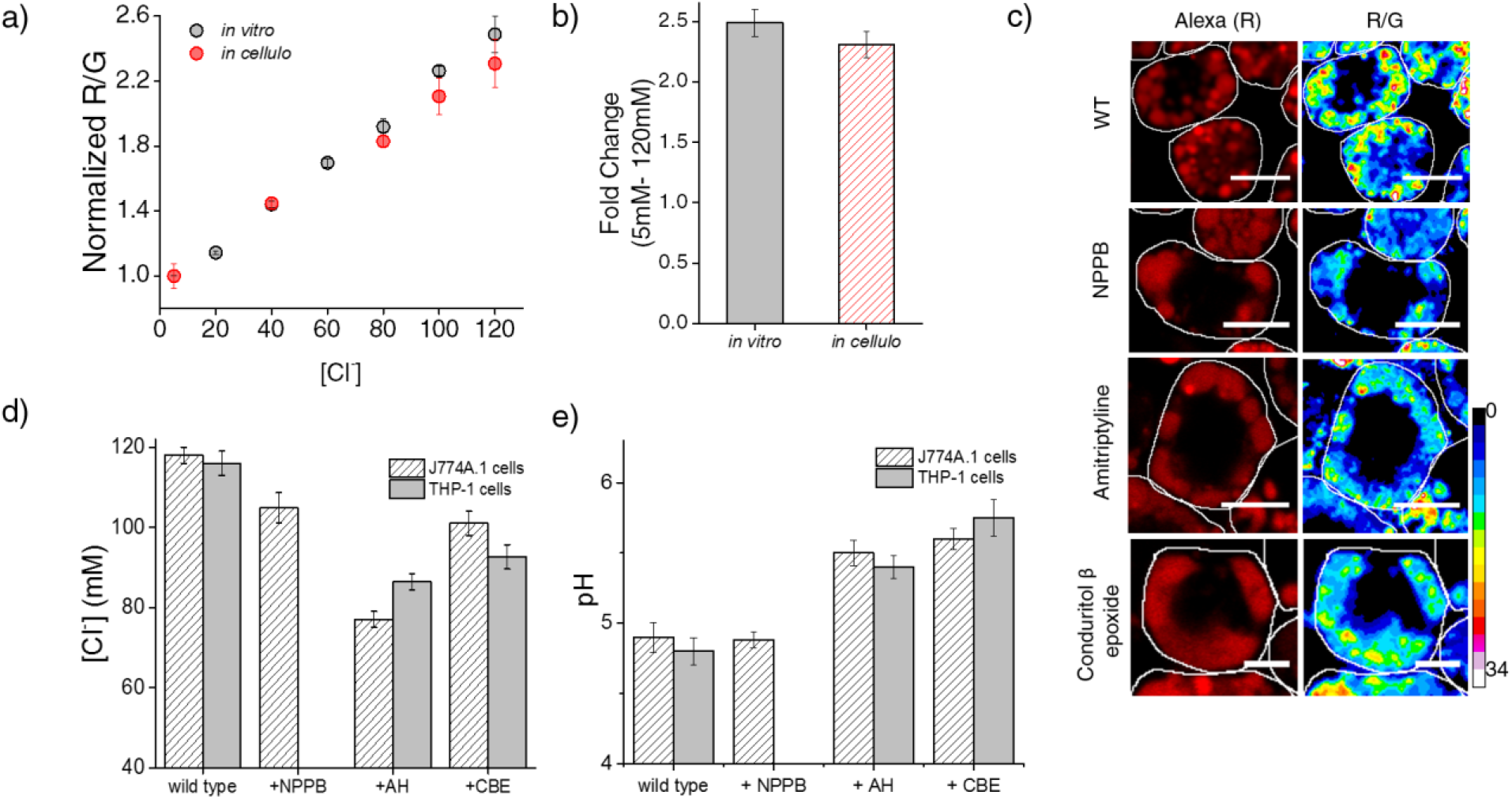
Lysosomal chloride is substantially depleted in mammalian cell culture models of lysosomal storage diseases. (a) Calibration profile of *Clensor* in cells (red) and *in vitro* (grey) showing normalized Alexa 647 (R) and BAC (G) intensity (R/G) ratios versus [Cl-]. Error bars indicate s.e.m. (n = 20 cells, ≥100 endosomes) (b) Fold change in R/G ratios of *Clensor in vitro* and in cells from 5 mM to 120 mM [Cl^−^] (c) Representative [Cl^−^] maps of *Clensor* in lysosomes of J774A.1 cells treated with the indicated lysosomal enzyme inhibitor. Images of the Alexa 647 (R) channel and pseudocolored R/G images are shown. Scalebar: 10 μm. (d) Bar graphs of lysosomal chloride values obtained in THP-1 and J774A.1 cells treated with the indicated inhibitors. NPPB (50 μM), Amitryptiline, AH (10 μM), Conduritol β-epoxide, CBE (400 μM) were used to model Niemann Pick A/B and Gaucher’s diseases in both cell types. Error bars indicate s.e.m. (n = 10 cells, ≥60 endosomes). (e) Bar graphs of lysosomal pH values obtained in THP-1 and J774A.1 cells treated with the indicated inhibitors. NPPB (50 μM), Amitryptiline, AH (10 μM), Conduritol β-epoxide, CBE (400 μM) were used to model Niemann Pick A/B and Gaucher’s diseases in both cell types. Error bars indicate s.e.m. (n = 10 cells, ≥50 endosomes).

Treating J774A.1 cells and THP-1 cells with a global chloride ion channel blocker, such as NPPB (5-Nitro-2-(3-phenylpropylamino) benzoic acid), lowered lysosomal chloride concentrations to 104 and 76 mM respectively, indicating that *Clensor* was capable of measuring pharmacologically induced lysosomal chloride changes if any, in these cells. In Gaucher’s cell culture models, murine and human cells showed a substantial decrease in lysosomal chloride to ~101 mM and ~92 mM respectively. This is a drop of 15–25 mM (13-21% change) chloride, as compared to a drop of ~10 μM in lysosomal proton concentrations. In Niemann-Pick A/B cell culture models, murine and human macrophages showed an even more dramatic decrease in lysosomal chloride to ~77 mM and ~86 mM respectively. This is also a substantial decrease of 30-40 mM (25-34% change) chloride, as compared to a drop of ~9 μM in lysosomal proton concentrations. On average in these four cell culture models, we find that the magnitude of chloride concentration decrease is at least 3 orders of magnitude greater than proton decrease, indicating that lysosome dysfunction is easily and sensitively reflected in its lumenal chloride concentrations. A Niemann Pick C cell culture model using the inhibitor U18666A recapitulated our findings in nematode models, where only lysosomal pH, but not Cl^−^, was altered (Figure 4 figure supplementary 5)

### High chloride regulates lysosome function in multiple ways

The ClC family protein CLC-7 is expressed mainly in the late endosomes and lysosomes (Graves et al., 2008; Jentsch, 2007). The loss of either ClC-7 or its β-subunit Ostm1 does not affect lysosomal pH in any way, yet leads to osteopetrosis, resulting in increased bone mass, and severe degeneration of the brain and retina (Lange et al., 2006). Along with our studies in nematodes, this reveals a role for high chloride in lysosome function that is beyond that of a mere counter-ion in the lysosome. We therefore probed whether it could indirectly affect lysosomal function by affecting lysosomal Ca^2+^ (Luzio et al., 2007; Rodríguez et al., 1997; Shen et al., 2012). We also considered the possibility that lysosomal chloride could exert a direct effect, where its reduction could impede the function of lysosomal enzymes thus affecting its degradative capacity (Baccino et al., 1975; Cigic and Pain, 1999; Maurus et al., 2005; Wartosch and Stauber, 2010) (Figure 5a).

**Figure 5:**
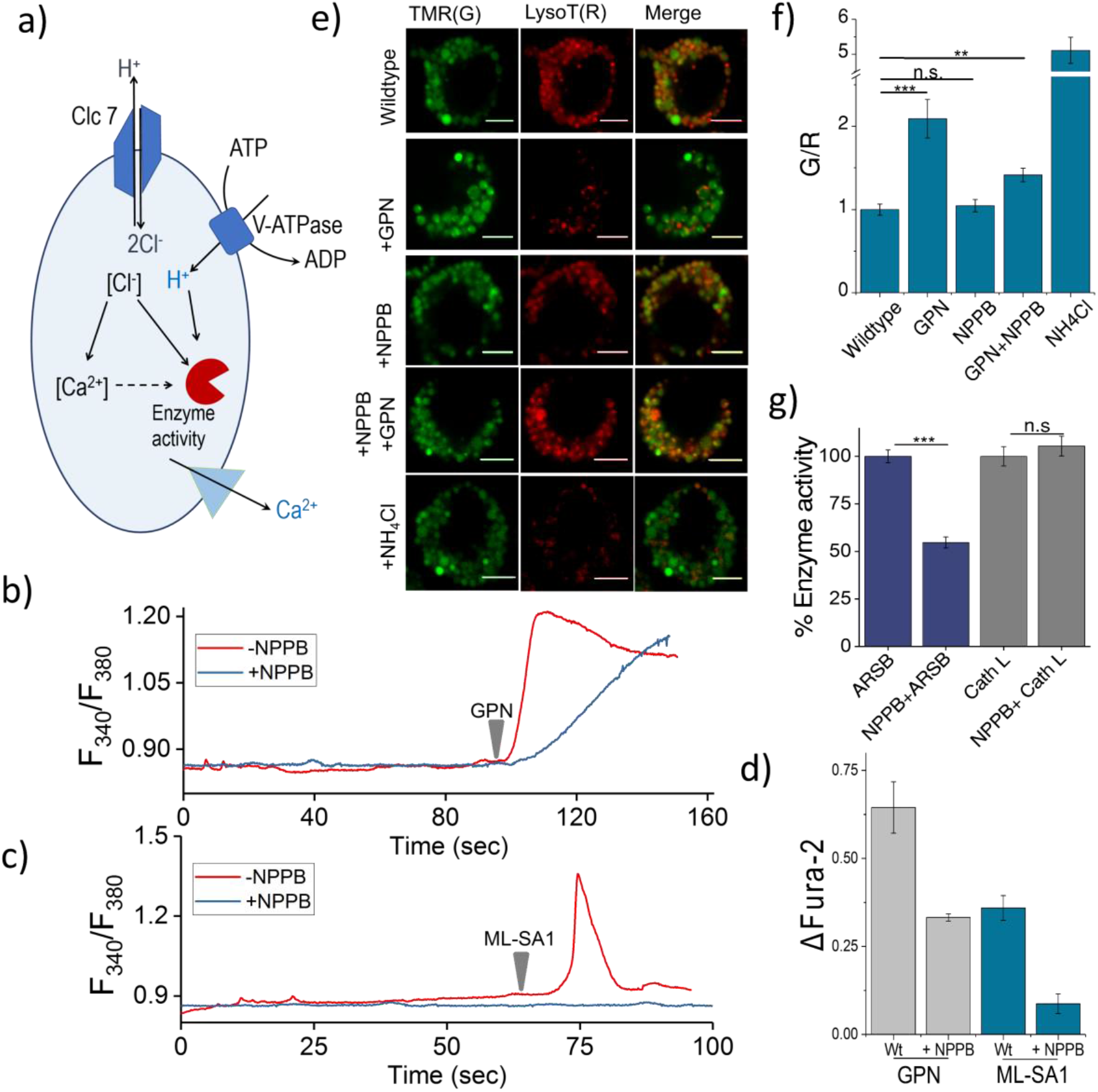
a) Schematic of potential roles for lysosomal Cl^−^. Chloride ions can regulate lysosomal Ca^2+^ and affect lysosomal enzyme function. b) Representative traces of GPN (400 μM) triggered lysosomal Ca^2+^ release in *J774A.1* cells ratiometrically imaged using Fura-2 (F_340_/F_380_) in the presence and absence of 50 μM NPPB. c) Representative traces of ML-SA1 (20 μM) triggered lysosomal Ca^2+^ release in *J774A.1* cells ratiometrically imaged using Fura-2 (F_340_/F_380_) in the presence and absence of 50 μM NPPB. d) Quantification of lysosomal data from e and f given by (F_t_-F_0_/F_0_) (ΔFura-2)for n = 15 cells. e) Representative images of lysosomes of *J774A.1* cells labelled with TMR dextran (TMR; G) and LysoTracker^®^ Red (LysoT; R) in the presence or absence of 50 μM NPPB, 200 μM GPN or 1 mM NH4Cl. Scale bar, 5 μm f) Quantification of LysoTracker^®^ release from b (n = 50 cells). g) Quantification of enzyme activity in J774A.1 cells in the absence and presence of 50 μM NPPB (n= 70 cells). Error bars indicate s.e.m. P values are as follows; ** = p<0.001, *** = p<0.0001, n.s = non significant.

Lysosomes are also intracellular Ca^2+^ stores with free Ca^2+^ ranging between ~400-600 μM (Christensen et al., 2002; Lloyd-Evans et al., 2008). The principal Ca^2+^ channel responsible for lysosomal Ca^2+^ release is Mucolipin TRP channel 1 (TRPML1). We therefore sought to estimate lysosomal Ca^2+^ by measuring Ca^2+^ that is released from the lysosome using two different triggers under conditions of normal and reduced lysosomal Cl^−^. Glycyl-L-phenylalanine 2-naphthylamide (GPN) is a substrate for Cathepsin C, which when added to cells, gets cleaved in the lysosome to release naphthylamine that is known to compromise the integrity of the lysosomal membrane, leading to a leakage of ions such as Ca^2+^ into the cytosol (Berg et al., 1994; Jadot et al., 1984; Morgan et al., 2011). This has been used to induce lysosomal Ca^2+^ release.

The cytosol of J774A.1 cells are labeled with ~3 μM Fura2-AM to ratiometrically image cytosolic Ca^2+^ elevation upon its release, if at all, from the lysosome. After addition of 400 μM GPN, cells were continuously imaged ratiometrically over 15–20 mins. Shortly after GPN addition, a burst of Ca^2+^ was observed in the cytosol, corresponding to released lysosomal Ca^2+^ (Figure 5b). When the same procedure was performed on cells that had been incubated with 50 μM NPPB that reduces lysosomal Cl-, the amount of lysosomal Ca^2+^ released was significantly reduced (Figure 5b–d) We then performed a second, more targeted way to release lysosomal Ca^2+^ into the cytosol, by using 20 μM ML-SA1 which specifically binds to and opens the TRPML1 channel on lysosomes (Shen et al., 2012). We found that when lysosomal Cl^-^ was reduced with NPPB, lysosomal Ca^2+^ release into the cytosol was near negligible (Figure 5c – d). Taken together this indicates that high lysosomal Cl^-^ is necessary for effective lysosomal Ca^2+^ release, possibly by affect lysosomal Ca^2+^ accumulation.

We next investigated whether reducing lysosomal chloride directly impacted the activity of any lysosomal enzymes. *In vitro* enzymology of Cathepsin C, a lysosome-resident serine protease has revealed that increasing Cl^−^ increased its enzymatic activity (Cigic and Pain, 1999; McDonald et al., 1966). Further, the crystal structure of Cathepsin C shows bound chloride ions close to the active site (Cigic and Pain, 1999; Turk et al., 2012). We therefore used GPN cleavage to probe Cathepsin C activity in the lysosome upon reducing Cl^−^ with NPPB. GPN cleavage by Cathepsin C releases naphthylamine which compromises lysosomal membrane integrity leading to proton leakage from the lysosome into the cytosol. This hypoacidifies the lysosomes resulting in reduced LysoTracker^®^ labeling as the labeling efficiency of the latter is directly proportional to compartment acidity.

Lysosomes are pre-labeled with TMR-Dextran, and LysoTracker^®^ intensities are normalized to the fluorescence intensity of TMR-Dextran, given as G/R. Hypoacidifying lysosomes by addition of 1 mM NH4Cl indeed reduced LysoTracker^®^ labeling, as expected (Figure 5e–f). A similar effect was also obtained upon GPN addition. The presence or absence of NPPB showed no change in LysoTracker^®^ labeling in cells (Figure 5e – f), indicating that NPPB by itself caused no alteration in lysosomal pH. However, when GPN was added to NPPB treated cells LysoTracker^®^ staining was remarkably well preserved (Figure 5e and f) indicating preservation of lysosomal membrane integrity because GPN was no longer effectively cleaved by Cathepsin C when lysosomal Cl^−^ was reduced. Unlike other cathepsins, Cathepsin C does not undergo autoactivation but requires processing by Cathepsin L and Cathepsin S to convert it into active Cathepsin C (Dahl et al., 2001). We measured the activity of the upstream cathepsins such as Cathepsin L using fluorogenic substrates in the presence and absence of NPPB (Figure 5g, Figure 5 figure supplementary 1). We observed no effect of chloride levels on Cathepsin L activity. This indicates that low Cathepsin C activity is not due to decreased amounts of mature Cathepsin C in the lysosome, but rather, reduced activity of mature Cathepsin C (Figure 5g, Figure 5 figure supplementary 1).

Based on reports suggesting that arylsulfatase B activity was also affected by low chloride (Wojczyk, 1986), we similarly investigated a fluorogenic substrate for arylsulfatase and found that NPPB treatment impeded arylsulfatase cleavage in the lysosome. Taken together, these results suggest that high lysosomal chloride is integral to the activity of key lysosomal enzymes and that reducing lysosomal chloride affects their function.

## Conclusions

The lysosome is the most acidic organelle within the cell. This likely confers on it a unique ionic microenvironment, reinforced by its high lumenal chloride, that is critical to its function (Xu and Ren, 2015). Using a DNA-based, fluorescent reporter called *Clensor* we have been able to create quantitative, spatial maps of chloride *in vivo* and measured lysosomal chloride. We show that, in *C. elegans*, lysosomes are highly enriched in chloride and that when lysosomal chloride is depleted, the degradative function of the lysosome is compromised. Intrigued by this finding, we explored the converse: whether lysosomes that had lost their degradative function – as seen in lysosomal storage disorders - showed lower lumenal chloride concentrations. In a host of *C. elegans* models for various lysosomal storage disorders, we found that this was indeed the case. In fact, the magnitude of change in chloride concentrations far outstrips the change in proton concentrations by at least four orders of magnitude.

To see whether chloride dysregulation correlated with lysosome dysfunction more broadly, we studied murine and human cell culture models of Gaucher’s disease, Niemann-Pick A/B disease and Niemann Pick C. We found that in mammalian cells too, lysosomes are particularly rich in chloride, surpassing even extracellular chloride levels. Importantly, chloride values in all the mammalian cell culture models revealed magnitudes of chloride dysregulation that were similar to that observed in *C. elegans*.

Our findings suggest more widespread and as yet unknown roles for the single most abundant, soluble physiological anion in regulating lysosome function. The ability to quantitate lysosomal chloride enables investigations into the broader mechanistic roles of chloride ions in regulating multiple functions performed by the lysosome. As such, given that chloride dysregulation shows a much higher dynamic range than hypoacidification, quantitative chloride imaging can provide a much more sensitive measure of lysosome dysfunction in model organisms as well as in cultured cells derived from blood samples that can be used in disease diagnoses and screening applications.

## Materials and Methods

### Reagents

All fluorescently labeled oligonucleotides were HPLC-purified and obtained from IBA-GmBh (Germany) and IDT (Coralville, IA, USA). Unlabeled oligonucleotides were purchased from IDT (Coralville, IA, USA). The peptide nucleic acids (PNA) oligomer, P was synthesized using standard solid phase Fmoc chemistry on Nova Syn^®^ TGA resin (Novabiochem, Germany) using analytical grade reagents (Applied Biosystems^®^, USA), purified by reverse phase HPLC (Shimadzu, Japan) as previously reported and stored at −20°C until further use (Prakash et al., 2016).

Bovine serum albumin (66 kDalton), nigericin, valinomycin, monensin, chloride ionophore I, IPTG, amitriptyline hydrochloride, 5-nitro-2-(3-phenylpropylamino) benzoic acid (NPPB) and conduritol β epoxide(CBE) were obtained from Sigma (USA). LysoTracker ^®^ Deep Red, TMR-Dextran (10kDa) and Oregon Green 488 maleimide was obtained from Molecular Probes, Invitrogen (USA). Lysosomal enzyme kits namely lysosomal sulfatase assay kit was purchased from Marker Gene (USA); Magic Red Cathepsin L assay kit from Immunochemistry Technologies. Gly-Phe β-naphthylamide was purchased from Santa Cruz Biotechnology (USA). All other reagents were purchased from Sigma-Aldrich (USA) unless otherwise specified. BSA was maleylated according to a previously published protocol (Haberland and Fogelman, 1985). Trizol was purchased from Invitrogen (U.S.A.).

### Sample preparation

All oligonucleotides were ethanol precipitated and quantified by their UV absorbance. For I-switch (I4^cLY^_A488/A647_) sample preparation, 5 μM of I4 and I4′ were mixed in equimolar ratios in 20 mM potassium phosphate buffer, pH 5.5 containing 100 mM KCl. The resulting solution was heated to 90 °C for 5 minutes, cooled to the room temperature at 5 °C/15 min and equilibrated at 4 °C overnight. Samples were diluted and used within 7 days of annealing. A sample of *Clensor* was similarly prepared using HPLC purified oligonucleotides and PNA oligomer at a final concentration of 10 μM by mixing D1, D2 and P (see Table S1 for sequence information) in equimolar ratios in 10 mM sodium phosphate buffer, pH 7.2 and annealed as described above. For I^mLy^, Oregon Green maleimide was first conjugated to the thiol labeled oligonucleotide (Hermanson, n.d.). Briefly, to 10μM thiol labelled oligonucleotide in HEPES pH 7.4, 500 μM of TCEP (tris-carboxyethylphosphine) was added to reduce the disulfide bonds. After 1 hour at RT, 50 μM Oregon Green Maleimide was added and the reaction was kept overnight at RT. The reaction mixture was purified using an Amicon cutoff membrane filter (3kDa, Millipore) to remove unreacted dye (Figure 4 figure supplementary 1). A sample of I^mLy^ was similarly prepared using HPLC purified oligonucleotides at a final concentration of 5μM by mixing I^mLY^ OG and I^mLY^_AT647_ (see Table S1 for sequence information) in equimolar ratios in 10 mM sodium phosphate buffer, pH 7.2 and annealed as described above. Prior to use, all buffer stock solutions were filtered using 0.22 μm disk filters (Millipore, Germany).

### *C. elegans* methods and strains

Standard methods were followed for the maintenance of *C. elegans*. Wild type strain used was the *C. elegans* isolate from Bristol (strain N2) (Brenner, 1974). Strains used in the study, provided by the Caenorhabditis Genetics Center (CGC), are *clh-6(ok791), unc-32(f131), unc-32(e189), ppk-3(n2835), gba-3(gk3287), ppt-1(gk139), and cln-3.2(gk41) I; cln-3.3(gk118) cln-3.1(pk479)*. Transgenics used in this study, also provided by the CGC, are *arIs37 [pmyo-3::ssGFP*], a transgenic strain that expresses ssGFP in the body wall muscles, which is secreted in the pseudocoelom and endocytosed by coelomocytes and *pwIs50 [lmp-1::GFP + Cb-unc-119(+)]*, a transgenic strain expressing GFP-tagged lysosomal marker LMP-1. Genes, for which mutants were unavailable, were knocked down using Ahringer library based RNAi methods (Kamath and Ahringer, 2003). The RNAi clones used were: *L4440* empty vector control, *ncr-1* (clone F02E8.6, Ahringer Library), *ostm1* (clone F42A8.3, Ahringer Library), *snx-3* (clone W06D4.5, Ahringer Library), *manba* (clone C33G3.4, Ahringer Library), *aman-1* (clone F55D10.1, Ahringer Library), *sul-3* (clone C54D2.4, Ahringer Library), gba-3 (clone F11E6.1, Ahringer Library) and *asm1* (clone B0252.2, Ahringer Library).

### Coelomocyte labeling experiments

Coelomocyte labeling and competition experiments were carried out with I4^cLY^_A647_, and *Clensor_A647_* as described previously by our lab (Surana et al., 2011). For microinjections, the samples were diluted to 100 nM using 1X Medium 1 (150 mM NaCl, 5 mM KCl, 1 mM CaCl_2_, 1 mM MgCl_2_, 20 mM HEPES, pH 7.2). Injections were performed, in the dorsal side in the pseudocoelom, just opposite to the vulva, of one-day old wild type hermaphrodites using an Olympus IX53 Simple Inverted Microscope (Olympus Corporation of the Americas, Center Valley, PA) equipped with 40X, 0.6 NA objective, and microinjection setup (Narishige, Japan). Injected worms were mounted on 2.0% agarose pad and anesthetized using 40 mM sodium azide in M9 buffer. In all cases labeling was checked after 1 h incubation at 22°C.

### Colocalization experiments

I4^cLY^_A647_ or *Clensor_A647_* sample was diluted to 500 nM using 1X Medium 1 and injected in 10 *arIs37 [pmyo-3::ssGFP]* hermaphrodites as described previously by our lab (Surana et al., 2011). Imaging and quantification of the number of coelomocytes labeled, after 1 hr of incubation, was carried out on the Leica TCS SP5 II STED laser scanning confocal microscope (Leica Microsystems, Inc., Buffalo Grove, IL) using an Argon ion laser for 488 nm excitation and He-Ne laser for 633 excitation with a set of dichroics, excitation, and emission filters suitable for each fluorophore. Cross talk and bleed-through were measured and found to be negligible between the GFP/Alexa 488/BAC channel and Alexa 647 channel.

### RNAi experiments

Bacteria from the Ahringer RNAi library expressing dsRNA against the relevant gene was fed to worms, and measurements were carried out in one-day old adults of the F1 progeny (Kamath and Ahringer, 2003). RNA knockdown was confirmed by probing mRNA levels of the candidate gene, assayed by RT-PCR. Briefly, total RNA was isolated using the Trizol-chloroform method; 2.5 μg of total RNA was converted to cDNA using oligo-dT primers. 5 μL of the RT reaction was used to set up a PCR using gene-specific primers. Actin mRNA was used as a control. PCR products were separated on a 1.5% agarose-TAE gel. Genes in this study that were knocked down by RNAi correspond to clh-6, ncr-1 and ostm-1 that showed expected 1.1 kb (clh-6); 1.1 kb (ncr-1); 0.9 kb (ostm-1) etc (Figure 1 figure supplementary 1).

### *In vitro* fluorescence measurements

Fluorescence spectra were measured on a FluoroMax-4 Scanning Spectrofluorometer (Horiba Scientific, Edison, NJ, USA) using previously established protocols(Modi et al., 2009; Saha et al., 2015). Briefly, I4^cLY^_A488/A647_ was diluted to 50 nM in 1X pH clamping buffer of desired pH for all *in vitro* fluorescence experiments. All samples were vortexed and equilibrated for 30 min at room temperature. The samples were excited at 488 nm and emission collected between 505 nm-750 nm. A calibration curve was obtained by plotting the ratio of donor emission intensity (D) at 520 nm and acceptor intensity (A) at 669 nm (for A488/A647) as a function of pH. Mean of D/A from three independent experiments and their s.e.m were plotted for each pH value. For *in vitro* calibration of I^mLy^, 50nM of the sensor is diluted into 1X pH clamping buffer of desired pH. Oregon Green and Atto 647N are excited at 490nm and 645nm respectively. Emission spectra for Oregon Green and Atto 647N were collected between 500–550 nm and 650–700 nm respectively. A calibration curve was obtained by plotting the ratio of Oregon Green (G) at 520 nm and Atto 647N (R) at 665 nm (for G/R) as a function of pH. Mean of G/R from three independent experiments and their s.e.m were plotted for each pH value.

For chloride measurements, 10 μM stock of *Clensor* was diluted to a final concentration of 200 nM using 10 mM sodium phosphate buffer, pH 7.2 and incubated for 30 min at room temperature prior to experiments. BAC and Alexa 647 were excited at 435 nm for BAC and 650 nm for Alexa 647 respectively. Emission spectra of BAC and Alexa 647 were collected between 495-550 nm and 650-700 nm respectively. In order to study the chloride sensitivity of *Clensor*, final chloride concentrations ranging between 5 mM to 80 mM were achieved by addition of microliter aliquots of 1 M stock of NaCl to 400 μL of sample. Emission intensity of BAC at 505 nm (G) was normalized to emission intensity of Alexa 647 at 670 nm (R). Fold change in R/G ratio was calculated from the ratio of R/G values at two specific values of [Cl^−^], either 5 mM and 80 mM or 5 mM and 120 mM as mentioned in the text.

### *In vivo* measurements of pH and chloride

pH clamping and measurement experiments were carried out with I4^cLY^_A488/A647_ as described by our lab previously(Modi et al., 2009; Surana et al., 2011). For microinjections, the I-switch sample was diluted to 500 nM using 1X Medium 1. Briefly, worms were incubated at 22°C for 1 hr post microinjection and then immersed in clamping buffers (120 mM KCl, 5 mM NaCl, 1 mM MgCl_2_, 1 mM CaCl_2_, 20 mM HEPES) of varying pH, containing 100 μM nigericin and 100 μM monensin. In order to facilitate entry of the buffer into the body, the cuticle was perforated at three regions of the body using a microinjection needle. After 75 mins incubation in the clamping buffer, coelomocytes were imaged using wide field microscopy. Three independent measurements, each with 10 worms, were made for each pH value.

Chloride clamping and measurements were carried out using *Clensor*. Worms were injected with 2 μM of *Clensor* and incubated at 22°C for 2 hrs. To obtain the chloride calibration profile, the worms were then immersed in the appropriate chloride clamping buffer containing a specific concentration of chloride, 100 μM nigericin, 100 μM valinomycin, 100 μM monensin and 10 μM chloride ionophore I for 45 mins at room temperature. Chloride calibration buffers containing different chloride concentrations were prepared by mixing the 1X chloride positive buffer (120 mM KCl, 20 mM NaCl, 1 mM CaCl_2_, 1 mM MgCl_2_, 20 mM HEPES, pH, 7.2) and 1X chloride negative buffer (120 mM KNO_3_, 20 mM NaNO_3_, 1 mM Ca(NO_3_)_2_, 1 mM Mg(NO_3_)_2_, 20 mM HEPES, pH 7.2) in different ratios.

For real-time lysosomal pH or chloride measurements, 10 hermaphrodites were injected with I4^cLY^_A488/A647_ or *Clensor* respectively and incubated at 22°C for one hour. Worms were then anaesthetized and imaged on a wide field inverted microscope for pH measurements and confocal microscope for chloride measurements.

### Cell culture methods and maintenance

Mouse alveolar macrophage J774A.1 cells were a kind gift from Prof Deborah Nelson, Department of Pharmacological and Physiological Sciences, the University of Chicago, cultured in Dulbecco’s Modified Eagle’s Medium/F-12 (1:1) (DMEM-F12) (Invitrogen Corporation,USA) containing 10% heat inactivated Fetal Bovine Serum (FBS) (Invitrogen Corporation, USA). THP-1 monocyte cell line was obtained from late Professor Janet Rowley’s Lab at the University of Chicago. Cells were cultured in RPMI 1640 containing 10 *%* heat-inactivated FBS, 10 mM HEPES, 2 mM glutamine, 100 U/ml penicillin, and 100 μg/ml streptomycin, and maintained at 37 °C under 5 % CO2. All reagents and medium were purchased from (Invitrogen Corporation,USA). THP-1 monocytic cells were differentiated into macrophages in 60 mm dishes containing 3 ml of the RPMI 1640 medium containing 10nM PMA over 48 h.

### *In cellulo* measurements pH and chloride

Chloride clamping and measurements were carried out using *Clensor* using a previously published protocol from our lab (Saha et al., 2015). J774A.1 and THP-1 cells were pulsed and chased with 2 μM of *Clensor*. Cells are then fixed with 200 μL 2.5% PFA for 2 min at room temperature, washed 3 times and retained in 1X PBS. To obtain the intracellular chloride calibration profile, perfusate and endosomal chloride concentrations were equalized by incubating the previously fixed cells in the appropriate chloride clamping buffer containing a specific concentration of chloride, 10 μM nigericin, 10 μM valinomycin, and 10 μM tributyltin chloride (TBT-Cl) for 1 h at room temperature.

Chloride calibration buffers containing different chloride concentrations were prepared by mixing the 1X chloride positive buffer (120 mM KCl, 20 mM NaCl, 1 mM CaCl_2_, 1 mM MgCl_2_, 20 mM HEPES, pH, 7.2) and 1X - chloride negative buffer (120 mM KNO_3_, 20 mM NaNO_3_, 1 mM Ca(NO_3_)_2_, 1 mM Mg(NO_3_)_2_, 20 mM HEPES, pH 7.2) in different ratios.

For real-time chloride measurements, cells are pulsed with 2 μM of *Clensor* followed by a 60-min chase. Cells are then washed with 1X PBS and imaged. To see whether *Clensor* can detect changes in Cl^−^ accumulation under perturbed conditions, we treated cells with 50 μM NPPB, which is a well-known non-specific Cl^−^ channel blocker. Cells were labeled with 2 μM *Clensor* for 30 mins and chased for 30 mins at 37°C. The cells were then chased for 30 mins in media containing 50 μM NPPB and then imaged.

To estimate the chloride accumulation in the lysosomes of Gaucher’s Disease in two cell models i.e, murine J774A.1 and human THP-1 cells, glucosylceramide storage was induced catalytically inactivating the enzyme acid β-glucosidase, using its well-known inhibitor conduritol β epoxide (CBE)(Grabowski et al., 1986; Schueler et al., 2004). These are both well-documented murine and human cell culture models of Gaucher’s disease. Macrophage cells were cultured with 300 μM CBE for 48 hours. Cells were then pulsed and chased with 2 μM *Clensor* as previously described.

To estimate chloride accumulation in the lysosomes of Niemann Pick A/B disease, the same murine and human cell lines were used, and the activity of acid sphingomyelinase (ASM) in these macrophage cell lines was inhibited using the well-known inhibitor, amitriptyline hydrochloride(Beckmann et al., 2014; Kornhuber et al., 2010). Cells were labeled with 2 μM *Clensor* for 30 mins and chased for 30 mins at 37°C. The cells were then chased for 30 mins in media containing 10 μM amitriptyline hydrochloride and then imaged.

*In cellulo* pH clamping and measurement experiments were carried out with I^mLy^ with modifications to protocols described by our lab previously(Modi et al., 2013, 2009). J774A.1 and THP-1 cells were pulsed and chased with 500 nM of I^mLy^. Cells are then fixed with 200 μL 2.5% PFA for 20 min at room temperature, washed 3 times and retained in 1X PBS. To obtain the intracellular pH calibration profile, perfusate and endosomal pH were equalized by incubating the previously fixed cells in the appropriate pH clamping buffer clamping buffers (120 mM KNO_3_, 5 mM NaNO_3_, 1 mM Mg(NO_3_)_2_, 1 mM Ca(NO_3_)_2_, 20 mM HEPES, MES and NaOAc) of varying pH, containing 25 μM nigericin and 25 μM monensin for 30 mins at room temperature. For real-time pH measurements, cells are pulsed with 500 nM of I^mLy^ followed by a 60-min chase. Cells are then washed with 1X PBS and imaged. pH measurements in the lysosomes of Gaucher’s Disease and of Niemann Pick A/B disease, in the two cell models i.e, murine J774A.1 and human THP-1 cells, were carried out similar to the protocol above using 500 nM of I^mLy^.

### Microscopy

Wide field microscopy was carried out on IX83 research inverted microscope (Olympus Corporation of the Americas, Center Valley, PA, USA) using a 60X, 1.42 NA, phase contrast oil immersion objective (PLAPON, Olympus Corporation of the Americas, Center Valley, PA, USA) and Evolve^®^ Delta 512 EMCCD camera (Photometrics, USA). Filter wheel, shutter and CCD camera were controlled using Metamorph Premier Ver 7.8.12.0 (Molecular Devices, LLC, USA), suitable for the fluorophores used. Images on the same day were acquired under the same acquisition settings. All the images were background subtracted taking mean intensity over an adjacent cell free area. Mean intensity in each endosome was measured in donor (D) and acceptor (A) channels. Alexa 488 channel images (D) were obtained using 480/20 band pass excitation filter, 520/40 band pass emission filter and a 89016-ET - FITC/Cy3/Cy5 dichroic filter. Alexa 647 channel images (A) were obtained using 640/30 band pass excitation filter, 705/72 band pass emission filter and 89016-ET - FITC/Cy3/Cy5 dichroic filter. For FRET channel images were obtained using the 480/20 band pass excitation filter, 705/72 band pass emission filter and 89016-ET - FITC/Cy3/Cy5 dichroic filter. Mean intensity in each endosome was measured in donor and acceptor channels. A ratio of donor to acceptor intensities (D/A) was obtained from these readings. Pseudocolor images were generated by calculating the D/A ratio per pixel. Confocal images were captured with a Leica TCS SP5 II STED laser scanning confocal microscope (Leica Microsystems, Inc., Buffalo Grove, IL, USA) equipped with 63X, 1.4 NA, oil immersion objective. Alexa 488 was excited using an Argon ion laser for 488 nm excitation, Alexa 647 using He-Ne laser for 633 excitation and BAC using Argon ion laser for 458 nm excitation with a set of dichroics, excitation, and emission filters suitable for each fluorophore.

Ratiometric calcium imaging of Fura-2 was carried out on an Olympus IX81 microscope equipped with a 40x objective, NA= 1.2. Excitation of Fura-2 is performed using 340/26 and 380/10 nm excitation filters, equipped with a 455nm dichroic mirror and a 535/40 nm emission filter. Exposure time was kept at 100 ms for all the imaging experiments to minimize phototoxicity.

### Image analysis

Images were analyzed with ImageJ ver 1.49 (NIH, USA). For pH measurements Alexa 488 and Alexa 647 images were overlapped using ImageJ and endosomes showing colocalization were selected for further analysis. Intensity in each endosome was measured in donor (D) and FRET (A) channels and recorded in an OriginPro Sr2 b9.2.272 (OriginLab Corporation, Northampton, MA, USA) file from which D/A ratio of each endosome was obtained. The mean D/A of each distribution were converted to pH according to the intracellular calibration curve. Data was represented as mean pH value ± standard error of the mean. Data for pH clamping experiments was analysed similarly.

For chloride measurements, regions of cells containing lysosomes in each Alexa 647 (R) image were identified and marked in the ROI plugin in ImageJ. The same regions were identified in the BAC (G) image recalling the ROIs and appropriate correction factor for chromatic aberration if necessary. After background subtraction, intensity for each endosome was measured and recorded in an Origin file. A ratio of R to G intensities (R/G) was obtained from these values by dividing the intensity of a given endosome in the R image with the corresponding intensity in the G image. For a given experiment, mean [Cl^−^] of an organelle population was determined by converting the mean R/G value of the distribution to [Cl^−^] values according to the intracellular calibration profile. Data was presented as mean of this mean [Cl^−^] value ± standard error of the mean. Data for chloride clamping experiments was analyzed similarly.

Colocalization of GFP and Alexa 647 was determined by counting the numbers of Alexa 647 positive puncta that colocalize with GFP and representing it as a Pearson’s correlation coefficient.

### Lysosomal labelling in coelomocytes

Temporal mapping of I-switch and *Clensor* was done in 10 worms of *pwIs50 [lmp-1::GFP + Cb-unc-119(+)]* as previously described by our lab (Surana et al., 2011). Briefly, worms were injected with 500 nM of I4^cLY^_A647_ or *Clensor_A647_*, incubated at 22°C for one hour, and then imaged using Leica TCS SP5 II STED laser scanning confocal microscope (Leica Microsystems, Inc., Buffalo Grove, IL, USA). Colocalization of GFP and I4^cLY^_A647_ or *Clensor_A647_* was determined by counting the numbers of Alexa647 positive puncta that colocalize with GFP positive puncta and expressing them as a percentage of the total number of Alexa 647 positive puncta. In order to confirm lysosomal labeling in a given genetic background, the same procedure was performed on the relevant mutant or RNAi knockdown in *pwIs50 [lmp-1::GFP + Cb-unc-119(+)]*.

### Statistics and general methods

All pH and chloride clamping experiments (Fig 1b, Fig 1 figure supplementary 2, Fig 4 figure supplementary 2) were performed in triplicates and the standard error of mean (s.e. m) values are plotted with the number of cells considered being mentioned in each legend. Experiment with murine macrophage, J774A.1 and THP-1 cells (Fig 4) has been performed in triplicates. Ratio of standard error of the mean is calculated for n = 20 cells and n= 10 cells and is plotted in Fig 4d and e respectively. All pH and chloride measurements in *C.elegans* of indicated genetic backgrounds (Fig 2c, 3c and Fig S5c) were carried out in n=10 worms and the standard error of mean (s.e.m) values are plotted with the number of cells considered being mentioned in each legend.

### DNA stability assay

Coelomocyte labeling for stability assay were carried out with I4^cLY^_A647_, and *Clensor_A647_*. For microinjections, the samples were diluted to 500 nM using 1X Medium 1 (150 mM NaCl, 5 mM KCl, 1 mM CaCl_2_, 1 mM MgCl_2_, 20 mM HEPES, pH 7.2). Post injection the worms are incubated at 22°C. After requisite time the injected worms are anesthetized in 40 mM sodium azide in M9 buffer and mounted on a glass slide containing 2% agarose pad. Worms were imaged using Olympus IX83 research inverted microscope (Olympus Corporation of the Americas, Center Valley, PA, USA). The average whole cell intensity in the Alexa 647 channel was plotted as a function of time (Figure 2; figure supplementary 2) J774A.1 and THP-1 macrophage cells labeling was carried out using 500 nM *Clensor_A647_* using 1X Medium 1 (150 mM NaCl, 5 mM KCl, 1 mM CaCl_2_, 1 mM MgCl_2_, 20 mM HEPES, pH 7.2). Cells were pulsed for 30 mins and then chased at 37°C for the indicated time points. The average whole cell intensity in the Alexa 647 channel was plotted as a function of time (Figure 4 figure supplementary 3).

### Fura-2AM Imaging

Cells were loaded with 3 μM Fura-2 AM in HBSS (137 mM NaCl, 5 mM KCl, 1.4 mM CaCl_2_, 1 mM MgCl_2_, 0.25 mM Na_2_HPO_4_, 0.44 mM KH_2_PO_4_, 4.2 mM NaHCO3and 10 mM glucose) at 37 °C for 60 min. Post incubation cells were washed with 1X PBS and then imaged in Low Ca^2+^ or Zero Ca^2+^ buffer (145 mM NaCl, 5 mM KCl, 3 mM MgCl_2_, 10 mM glucose, 1 mM EGTA, and 20 mM HEPES (pH 7.4)). Ca^2+^ concentration in the nominally free Ca^2+^ solution is estimated to be 1–10 μM. With 1 mM EGTA, the free Ca^2+^ concentration is estimated to be <10 nM based on the Maxchelator software (http://maxchelator.stanford.edu/). Florescence was recorded using two different wavelengths (340 and 380 nm) and the ratio (F_340_/F_380_) was used to calculate changes in intracellular [Ca^2+^].

### Enzyme activity assays

Enzyme activity assays were performed in J774A.1 cells. For Cathepsin C enzyme activity; we used Gly-Phe β-naphthylamide as a substrate. Lysosomes of J774A.1 cells were pre-labeled with TMR-dextran (0.5 mg/mL; G) for 1 hour and chased in complete medium for 16 hours at 37°C. The cells were then labeled with 50 nM LysoTracker^®^ in complete medium for 30 mins at 37°C. 50 μM NPPB or 200 μM GPN were then added to the cells and incubated for 30 mins at 37°C. The cells then washed and imaged in HBSS buffer containing either NPPB or GPN. The whole cell intensity ratio (G/R) was plotted to quantify the level of LysoTracker^®^ labelling of the endosomes. For Cathepsin L and Aryl Sulfatase enzyme activity Magic Red Cathepsin L assay kit (Immunochemistry Technologies) and Lysosomal sulfatase assay kit (Marker Gene) were used. The experiment was performed using the manufacture’s protocol. Briefly, cells were incubated with 1X Magic Red Cathepsin L assay probe or 200 μM Lysosomal sulfatase assay probe for 4 hours in complete medium. The cells were then labelled with 10 μM Hoechst stain for 10 mins at 37°C after which the cells were washed and imaged. For low chloride containing dishes; cells were preincubated with 50 μM NPPB before the addition of the enzyme probes. The ratio of enzyme substrate whole cell intensity to that of DAPI was used to quantify enzyme activity.

## Acknowledgments

The authors thank Professors John Kuriyan, Susan Cotman, Drs A. H. Rahmathullah and D. McEwan for critical comments and valuable suggestions. The authors thank the Integrated Light Microscopy facility at the University of Chicago, the *C. elegans* Genetic Center (CGC) for strains, Koushika, S., Glotzer, M., and Ausbel, F. for Arhinger Library RNAi clones. Data described in the paper are presented in the main text and the supplementary materials. This work was supported by the National Center for Advancing Translational Sciences of the National Institutes of Health through grant no: UL1 TR000430 and U Chicago startup funds to YK. YK is a Brain Research Foundation Fellow.

---

**Figure 1 figure supplementary 1:**
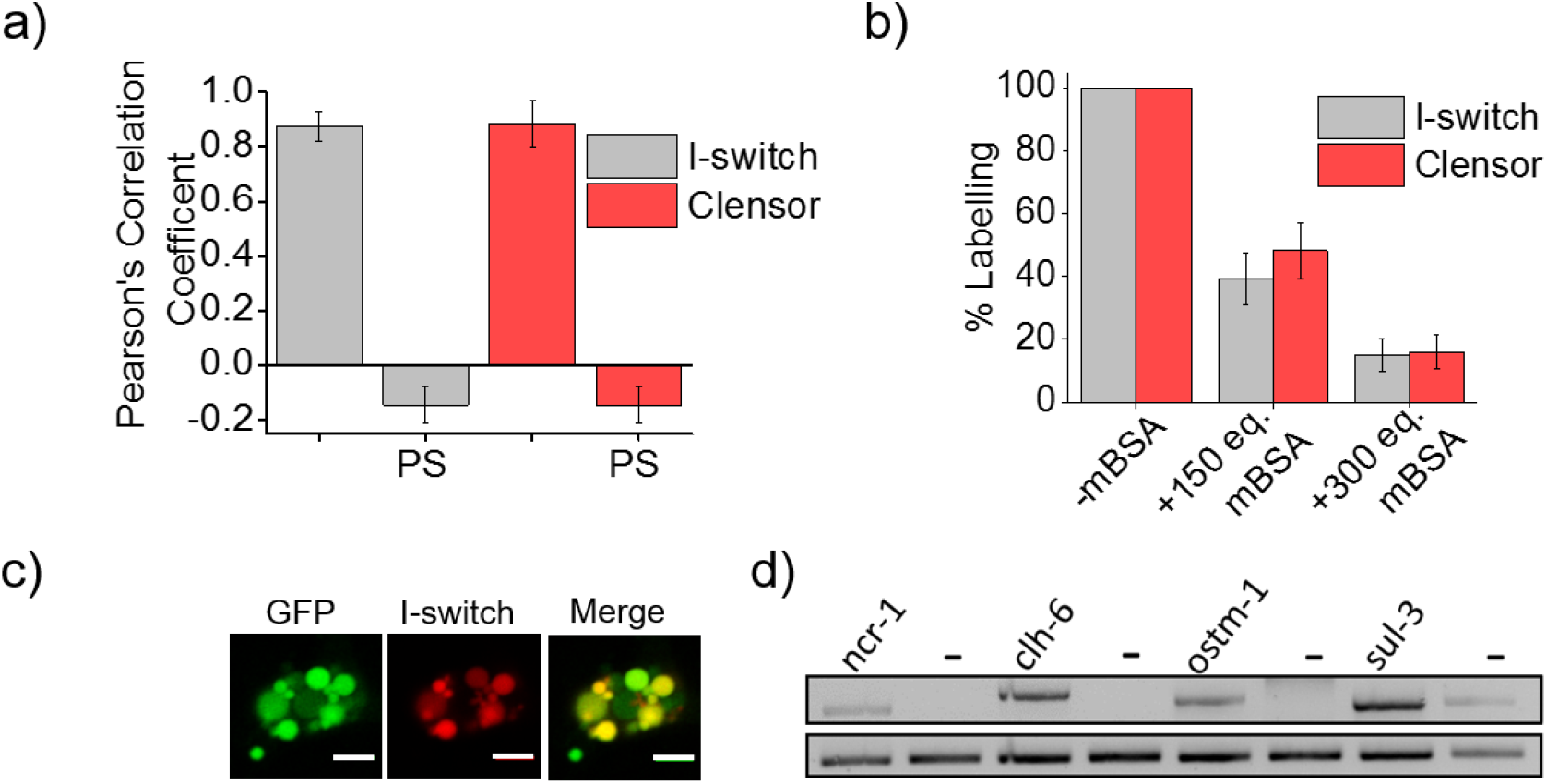
(a) Quantification of co-localization between the DNA nanodevices and GFP in arIs37 worms. Mean of n=10 cells. (b) Coelomocyte labeling efficiency with *I-switch* (I4_A647_) and *Clensor_A647_* in the absence or presence of 150 or 300 equivalents excess of maleylated bovine serum albumin (mBSA) Error bars indicate s.e.m. (n = 10 worms) (c) Images showing colocalization of *Clensor_A647_* or *I-switch* (red) 1 h post injection in the pseudocoelom of arIs37 [pmyo-3::ssGFP] worms (green). Scale bar: 5 μm. (d) RT-PCR analysis of total RNA isolated from *C. elegans* pre- and post-RNAi. Lanes correspond to PCR-amplified cDNA of the indicated gene product isolated from wild type without RNAi treatment (denoted by gene name) and dsRNA-fed worms (denoted as “-ncr-1, chl6, ostm1 and sul3”).

**Figure 1 figure supplementary 2:**
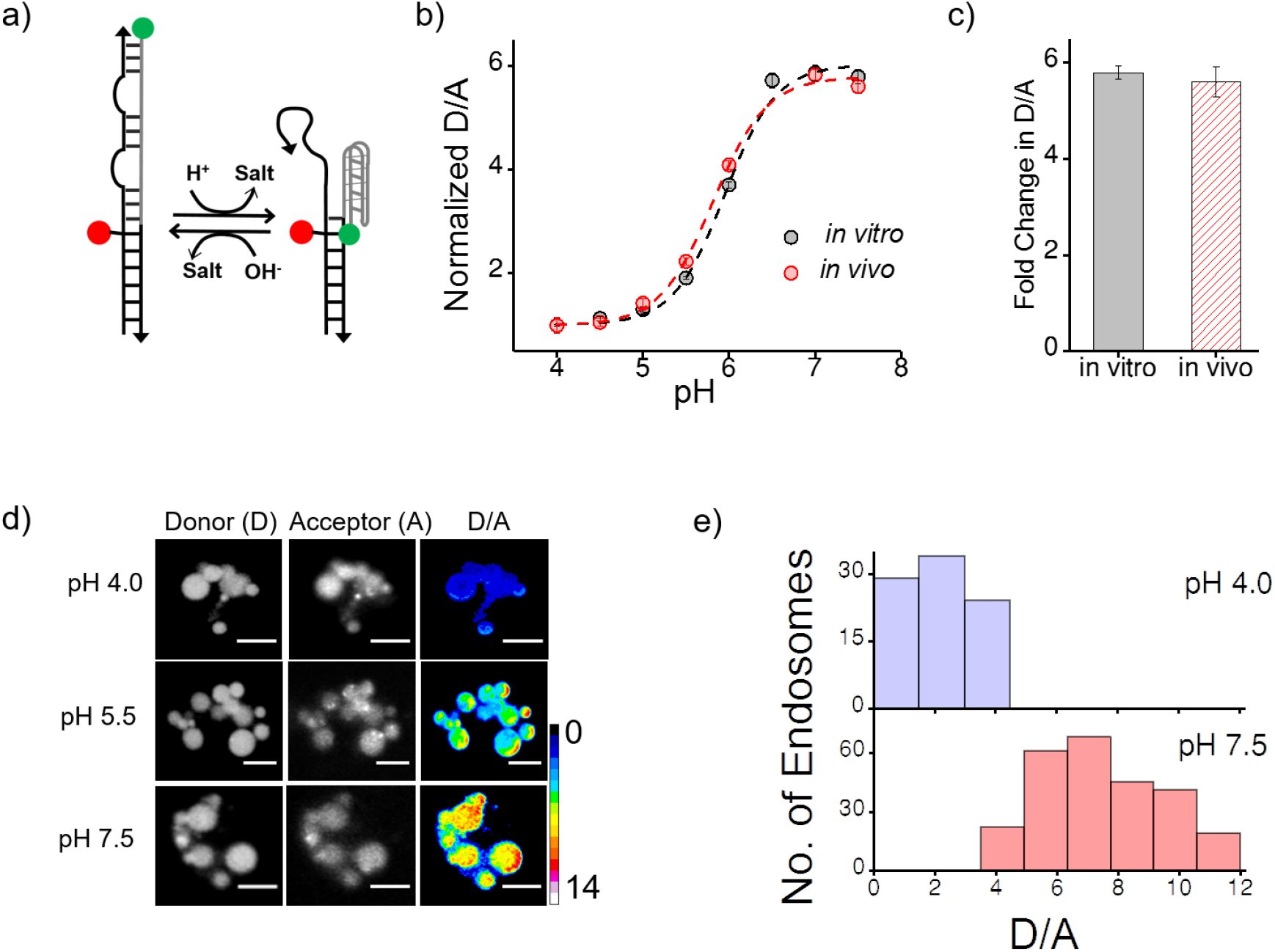
(a) Schematic of a DNA nanodevice, the *I-switch*, that functions as a fluorescent pH reporter based on a pH triggered conformational change that is transduced to photonic changes driven by differential fluorescent resonance energy transfer between donor (green) and acceptor (red) fluorophores (b) pH calibration curve of I4^cLY^_A488/A647_ *in vivo* (black) and *in vitro* (red) showing normalized D/A ratios versus pH. Error bars indicate s.e.m. (n = 15 cells, ≥60 endosomes) (c) *in vitro* and *in vivo* fold change in D/A ratios of I4^cLY^_A488/A647_ from pH 4.0 to pH 7.5. (d) Representative images of coelomocytes labelled with I4^cLY^_A488/A647_, clamped at the indicated pH. Images are acquired in donor channel (D), acceptor channel (A) from which the respective pseudocolored D/A image is obtained. Scale bar, 5 μm. (e) Histograms showing typical spread of D/A ratios of endosomes clamped at pH 4 (lavender) and pH 7.5 (salmon);(*n* = 10 cells, ≥100 endosomes)

We use the I switch for pH measurements in the lysosomes of coelomocytes (Fig 1 figure supplementary 2). We first validate the sensor *in vivo*. For this we first generate an *in vivo* calibration profile for the sensor using previous methods standardized by our lab (Surana et al., 2011). Briefly worms are injected with 500nM I-switch and were placed in clamping buffers of varying pH 1h post injection. Post 75 mins incubation, worms were imaged in the Donor Channel (D) and Acceptor Channel (A) (Fig. 1 figure supplementary 2d). On plotting the mean D/A against varying pH we generate a calibration curve which shows that the sensor’s *in vivo* performance is similar to the *in vitro* profile (Fig. 1 figure supplementary 2b and c).

**Figure 1 figure supplementary 3:**
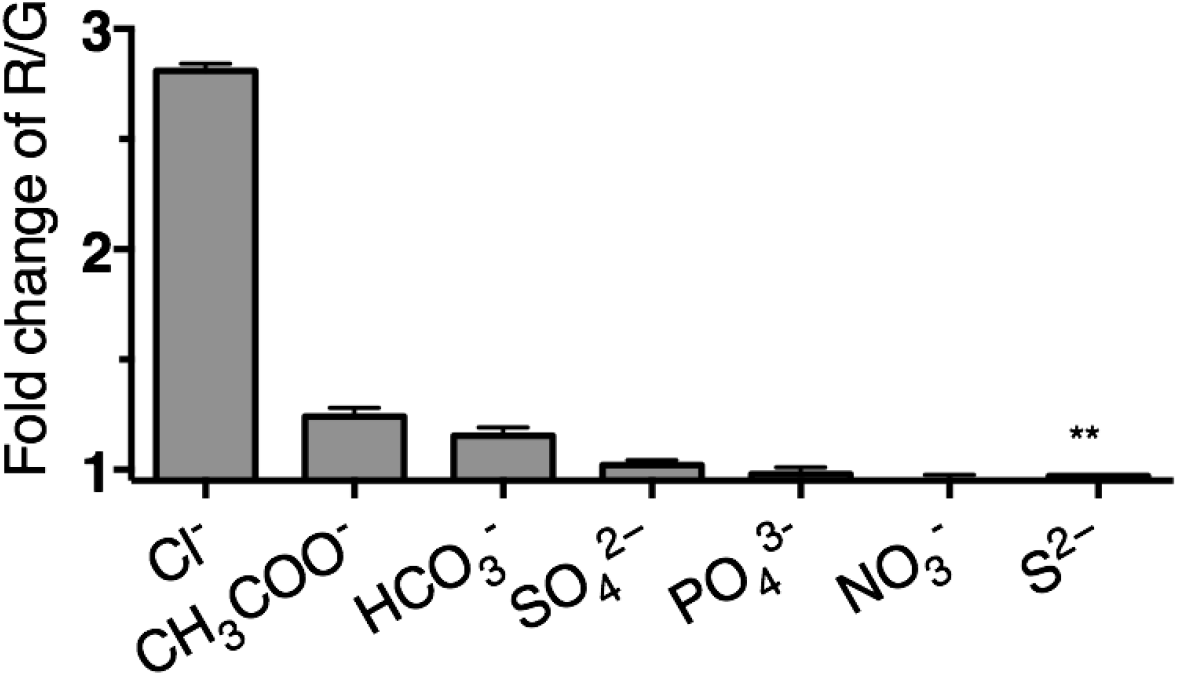
Selectivity of *Clensor* (200 nM) in terms of its fold change in R/G from ~ 0 mM to 100 mM of each indicated anion unless otherwise indicated. (* ~ 0 mM to 10 mM).

Since Cl^−^ channels are known to exhibit poor anion selectivity, we investigate the selectivity of *Clensor* to demonstrate the signal change in cell is attributed by the change of chloride ions level. The selectivity of *Clensor* to various anions in the form of their sodium salts. This reveals that the sensitivity of *Clensor* to diverse biologically abundant anions including NO_3_^−^ and PO_4_^3−^ is negligible.

**Figure 2 figure supplementary 1:**
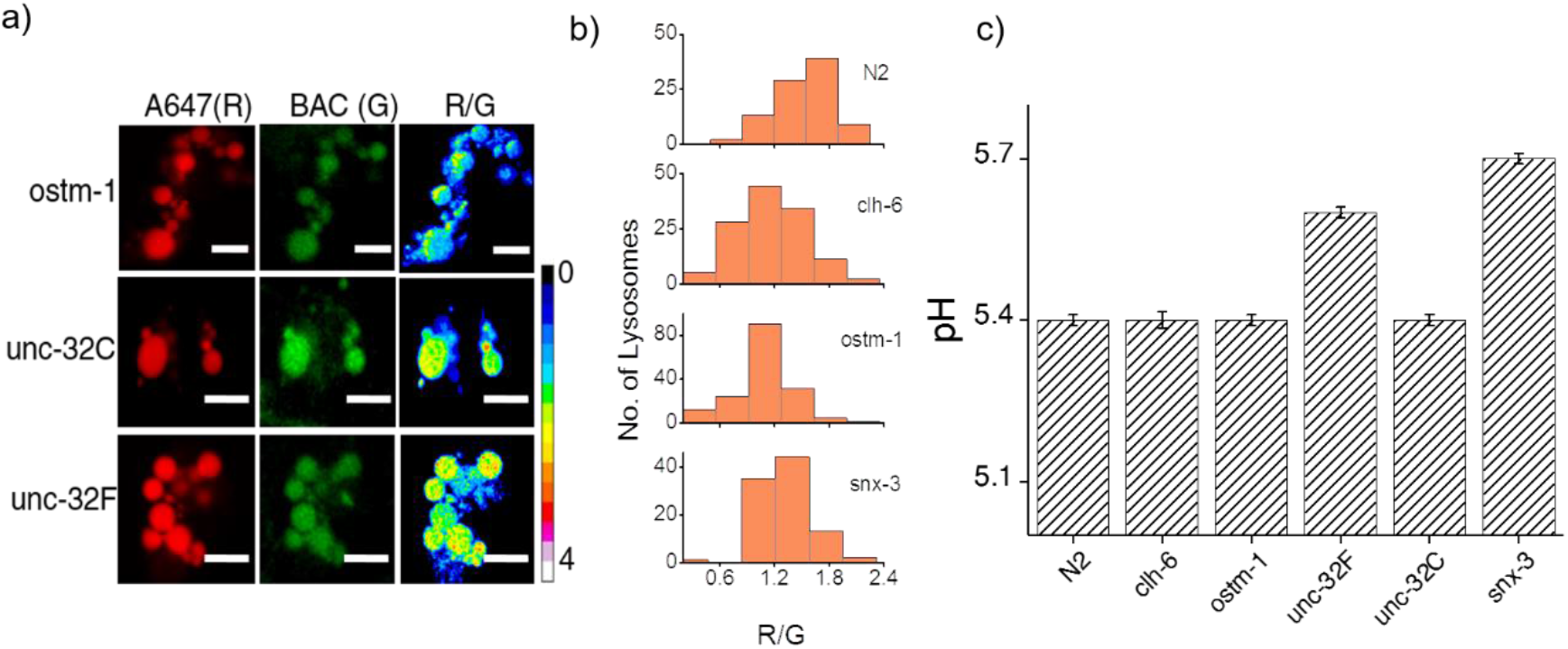
(a) Representative images of coelomocyte lysosomes labeled with *Clensor* one hour post injection, in the indicated genetic backgrounds acquired in the Alexa 647 (R) and BAC (G) channels and the corresponding pseudocolored R/G images. Scale bar, 5 μm. (b) Histograms comparing the spread of R/G in coelomocytes in different RNAi background. (n=10 cells; >100 lysosomes). (c)) Lysosomal pH measured using I4^cLY^_A488/A647_ in the indicated genetic backgrounds (n = 10 worms, ≥100 lysosomes).

Figure 2 figure supplementary 1a shows representative images of chloride measurements in lysosomes of coelomocytes of worms where various genes related to osteopetrosis are either knocked out or knocked downs. Fig. 2 figure supplementary 1b shows a histogram comparing the spread of R/G values reporting lysosomal chloride in the indicated genetic backgrounds. We observe that the R/G values are lower in the case of *clh-6* mutant and *ostm-1* RNAi whereas that of *snx-3* is similar to wild type.

**Figure 2 figure supplementary 2:**
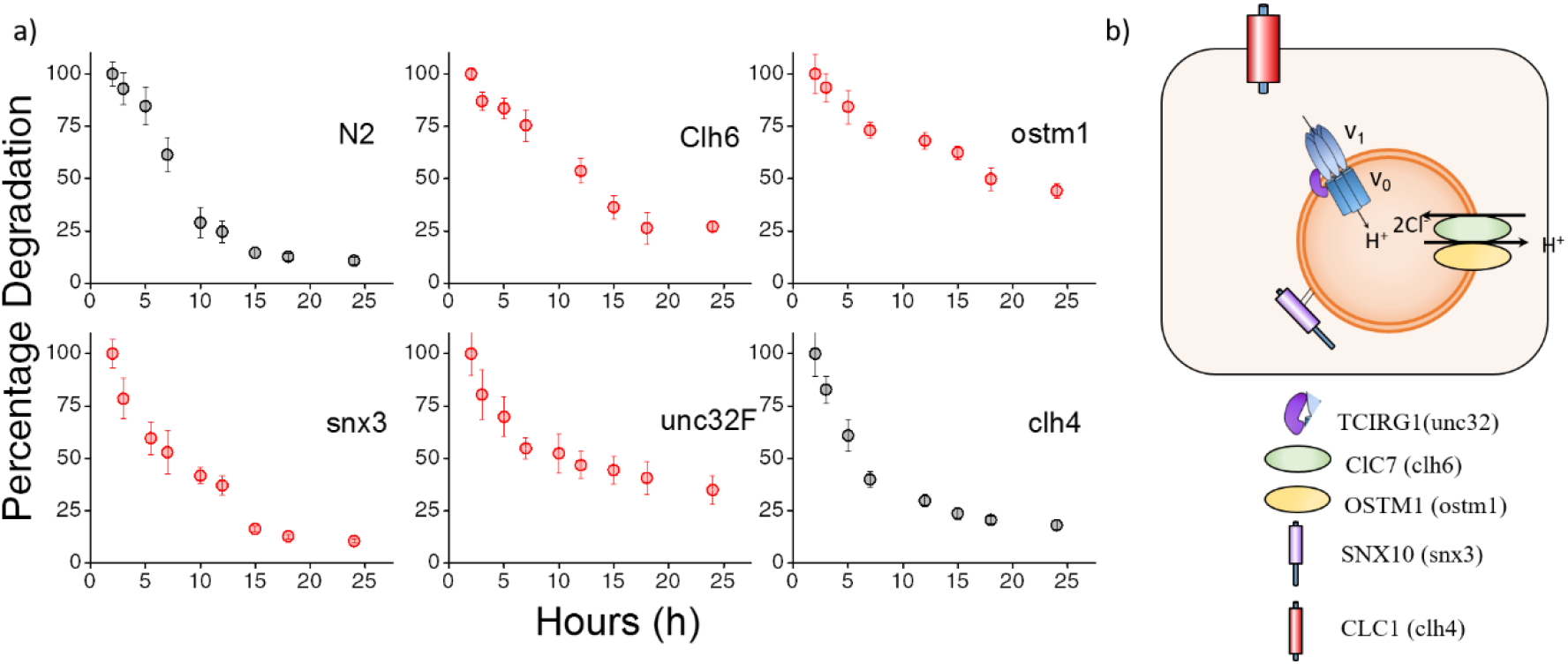
(a) Plots showing mean whole cell intensity per coelomocyte of I4_A647_, as a function of time post-injection in indicated genetic backgrounds. Error bars indicate s.e.m. (n ≥ 10 cells). (b) Schematic depicting protein players involved in autosomal recessive osteopetrosis and clh4, a plasma membrane resident chloride channel.

The above figure shows the stability of I switch in the lysosomes of the coelomocytes. Worms were injected with *I4*_A647_ and the number of labeled coelomocytes were plotted as a function of time. This gives an exponential decay profile, with progressively more coelomocytes losing their labeling as the DNA device is degraded in the coelomocyte. An exponential is fitted that gives the half-life of the device. In N2 worms, the half-life of the DNA reporters is ~ 6.5 hours, with maximal labelling occurring at 2 hours post injection.

**Figure 3 figure supplementary 1:**
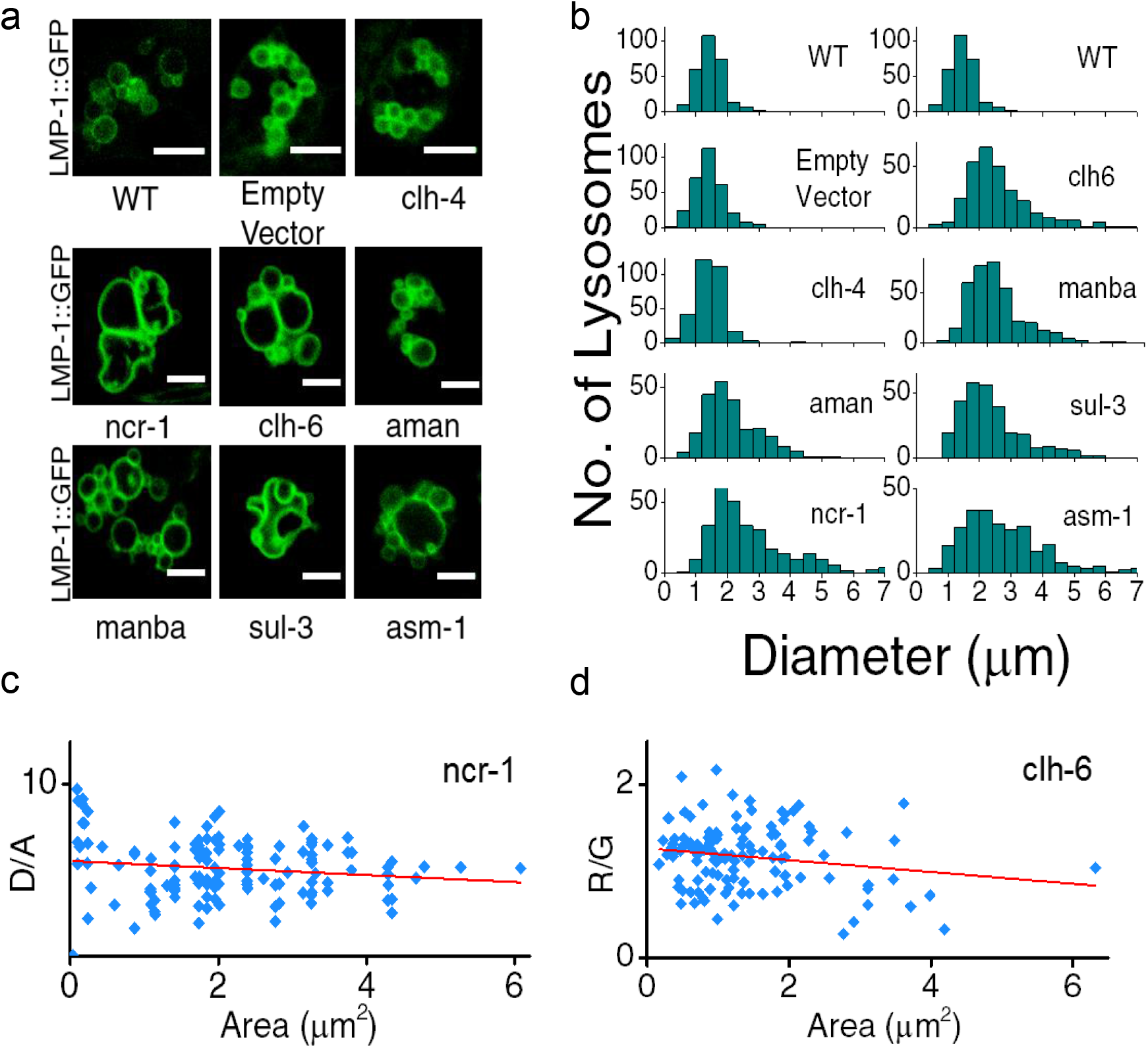
(a) Representative images of LMP-1::GFP marked coelomocytes in the background of indicated RNAi. Scale bar: 5 μm. (b) Histograms comparing the spread in size of LMP-1::GFP positive vesicles in coelomocytes in the indicated RNAi background (n = 20 cells). (c) D/A values obtained for individual pH measurements in lysosomes as a function of area of each vesicle in the indicated genetic background. The linear regression is shown in red. The coefficient of regression (p = −0.14) shows no correlation (n >100 lysosomes). (d) R/G values obtained for individual Cl^−^ measurement in lysosomes as a function of area of each vesicle in the indicated genetic background. The linear regression line is shown in red. The coefficient of regression (p= −0.18) shows no correlation (n >100 lysosomes).

A key phenotype observed in cells derived from patients suffering from lysosomal storage disorders (LSD) is the presence of enlarged lysosomes (Filocamo and Morrone, 2011; Platt et al., 2012). LSD-related gene knockdowns in worms show subtle to no observable whole organismal phenotypes (de Voer et al., 2008). However, on knocking down various LSD-related genes in the background of LMP-1::GFP worms we observed that the morphology of LMP-1 positive vesicles were altered and that the coelomocytes contained enlarged lysosomes, some as large as 8 microns in diameter (Fig. 3 figure supplementary 1). Fig 3 figure supplementary 1b represents a plot of the diameter distribution for LMP-1 positive vesicles under each indicated genetic background. Worms with LSD-related gene knockdowns show a broader distribution of vesicle sizes compared to a more tightly regulated size distribution in wild type nematodes or non-LSD related mutants. E.g., when *clh-4*, a plasma membrane resident chloride channel was knocked down, lysosomal morphology is not affected. We carried out pH and chloride measurements for worms from various genetic backgrounds and checked for correlations between lysosomal D/A (or R/G) and lysosome size. On plotting D/A values obtained from pH measurements in *ncr-1* RNAi worms, against the area of each vesicle; we observe no correlation between the two parameters (Fig. 3 figure supplementary 1c). A similar observation is seen in the case of chloride R/G measurements (Fig. 3 figure supplementary 1d). Thus assaying only for lysosome size shows no correlation with lysosome functionality. However lumenal chloride concentration is the best correlate of lysosome dysfunction, irrespective of size.

**Figure 3 figure supplementary 2:**
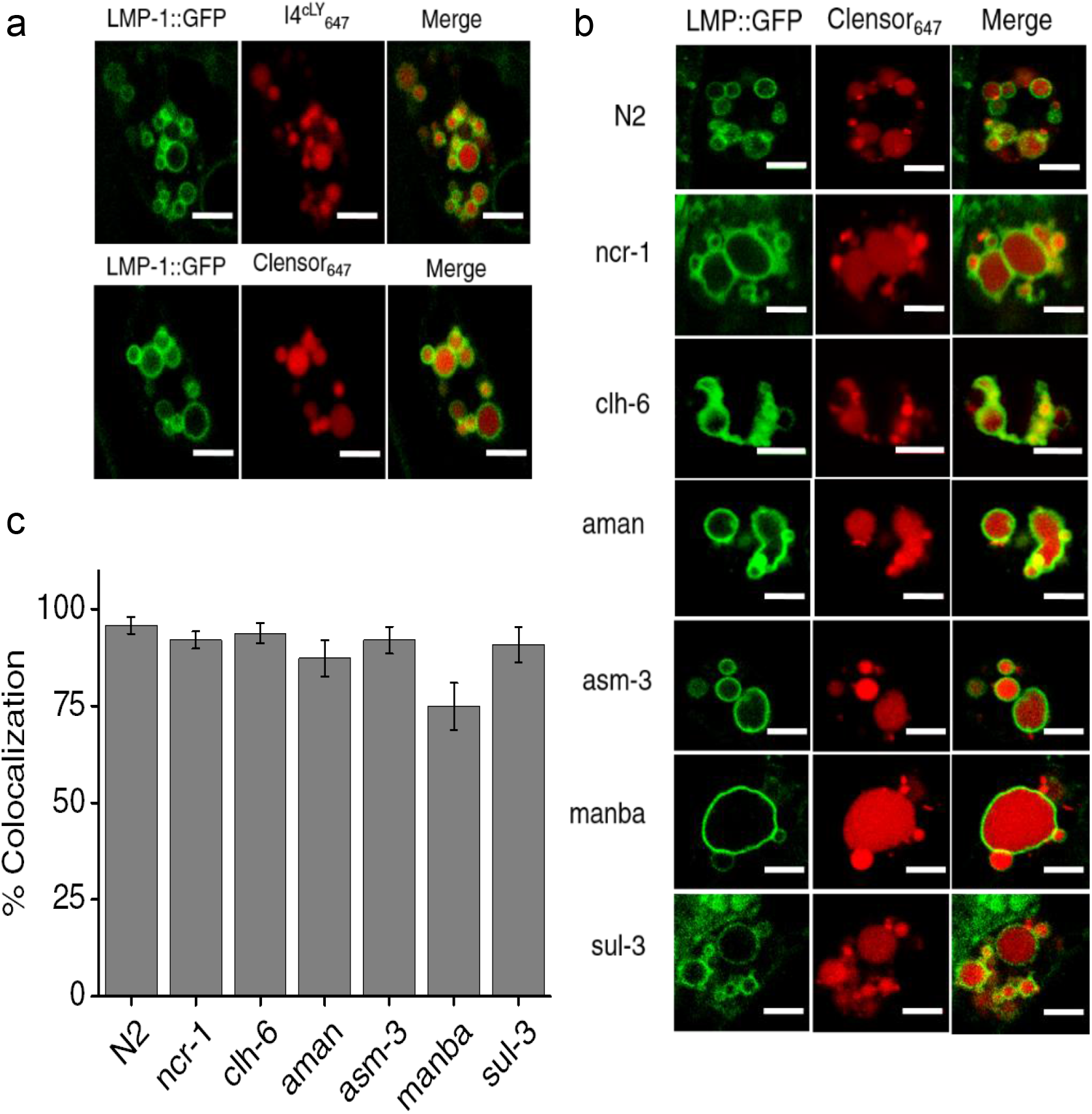
(a) Worms expressing LMP-1::GFP in coelomocytes were injected with I4^cLY^_A647_ or *Clensor_A647_* positive vesicles (red) and show maximal colocalization with LMP-1::GFP vesicles (green) at 60 min. Scale bar: 5 μm (b) Representative images of worms expressing LMP-1::GFP (green) in the background of various indicated RNAi’s, which were injected with *Clensor_A647_* (red) and imaged 60 mins post-injection. Scale bar: 5 μm. (c) Quantification of colocalization between the *Clensor_A647_* and GFP in arIs37 worms. Mean of n=10 cells

To check whether our DNA nanodevices can mark the lysosomes of coelomocytes in wild type worms, we injected 500 nM of I4^cLY^_A647_ or *Clensor_A647_* into worms containing LMP1::GFP marker (Fig. 3 figure supplementary 2a). The worms were imaged 1 h post injection in the GFP channel (green) and in Alexa 647 channel (red) to visualize the lysosomal marker and DNA reporter respectively. Merged images show colocalization of DNA devices with the lysosomal marker, similar to previous studies (Surana et al., 2011). We then proceeded to validate the devices in the lysosomes in coelomocytes of various LSD-related genes knocked down worms. LMP1::GFP positive worms that were RNAi-ed for indicated genes were injected with *Clensor_A647_* and imaged 1 h post injection (Fig. 3 figure supplementary 2b). We observe that, in worms, *Clensor* reliably marks the lysosomes in all LSD-related gene knockdowns with over 74% colocalization (Fig. 3 figure supplementary 2c).

**Figure 3 figure supplementary 4:**
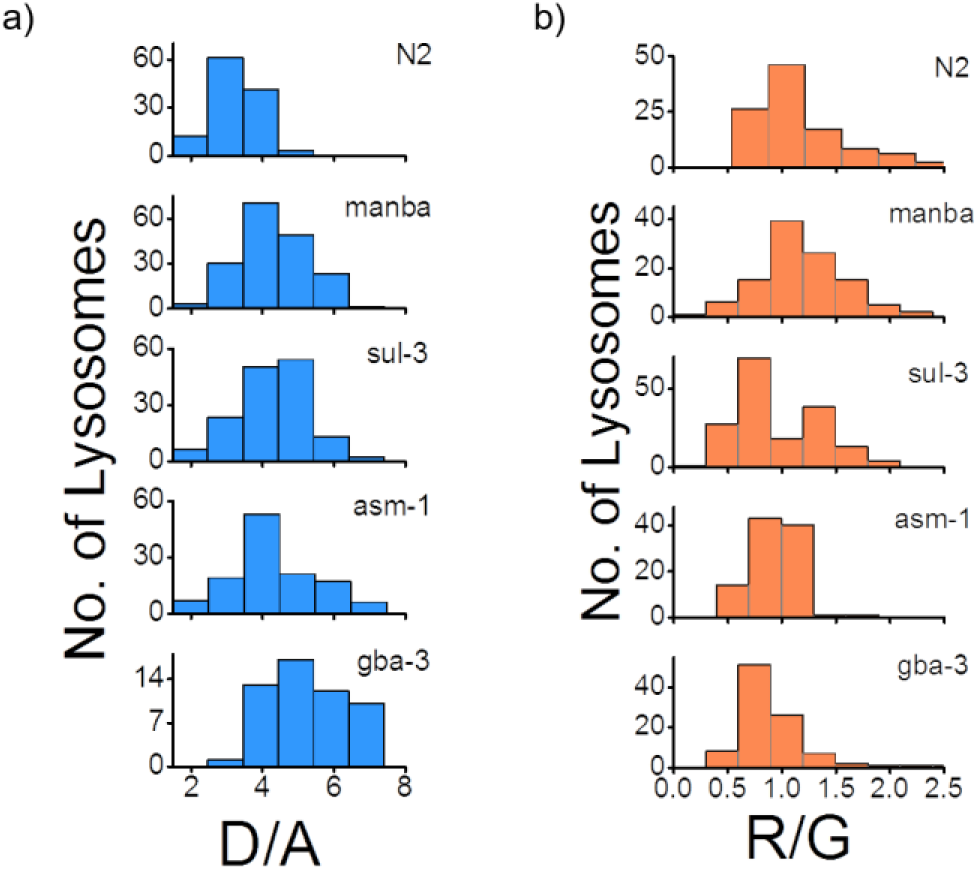
(a) Histograms comparing the spread of D/A in coelomocytes in different RNAi background. (n = 10 cells; >100 lysosomes). (b) Histograms comparing the spread of R/G in coelomocytes in different RNAi background. (n = 10 cells; >100 lysosomes).

**Figure 4 figure supplementary 1:**
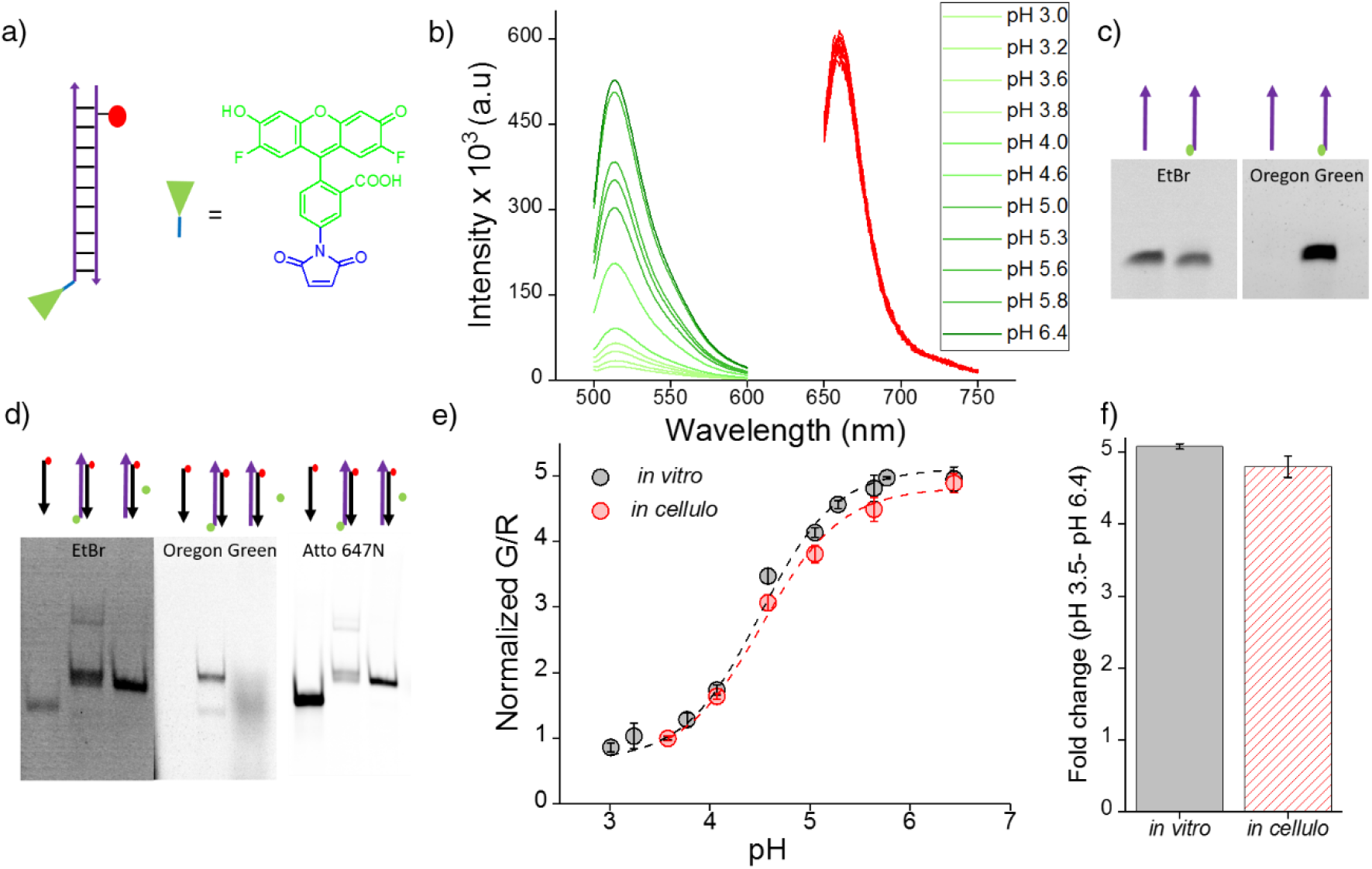
(a) Structure and schematic of I^mLy^ (b) Fluorescence emission spectra of I^mLy^ at the indicated pH obtained using λ_Ex_ OG = 494 nm (green) and λ_Ex_ Atto 647N = 650 nm (red). (c) Representative 12% denaturing PAGE showing conjugation of Oregon green (green circle) to thiol labelled DNA (purple arrow) (d) Representative 20% PAGE showing formation of I^mLy^ (e) pH calibration curve of I^mLy^ *in cellulo* (red) and *in vitro* (black) showing normalized G/R ratios versus pH. Error bars indicate s.e.m. (n = 15 cells, ≥60 endosomes) (f) *in vitro* and *in vivo* fold change in G/R ratios of I^mLy^ from 3.5 to pH 6.4.

To create a pH sensor for mammalian lysosomes we conjugated a pH sensitive dye – Oregon green (OG) which pKa is 4.9 (green chemical structure) to thiol labelled DNA using a maleimide linker (blue chemical structure). The complementary strand contains a normalizing dye, Atto 647N (red circle). The conjugation of OG was confirmed by 12% denaturing PAGE (Fig. 4 figure supplementary 1). Furthermore, the formation of I^mLy^ was confirmed by a gel mobility shift assay using native polyacrylamide gel electrophoresis. We annealed 10 μM of both components in equimolar ratios in 10 mM sodium phosphate buffer, pH 7.4 and investigated the sample by PAGE. The formation of I^mLy^ was revealed by its lower electrophoretic mobility with respect to its component single strands (Fig. 4 figure supplementary 1d). The fluorescence of OG increases with the pH while the fluorescence of normalizing dye Atto 647N remains unchanged (Fig. 4 figure supplementary 1b). Furthermore, the *in cellulo* calibration were well matched with their *in vitro* calibration profiles (Fig. 4 figure supplementary 1e–f), indicating that both sensor integrity and performance were quantitatively preserved at the time of making lysosomal pH a measurements in these cells.

**Figure 4 figure supplementary 2:**
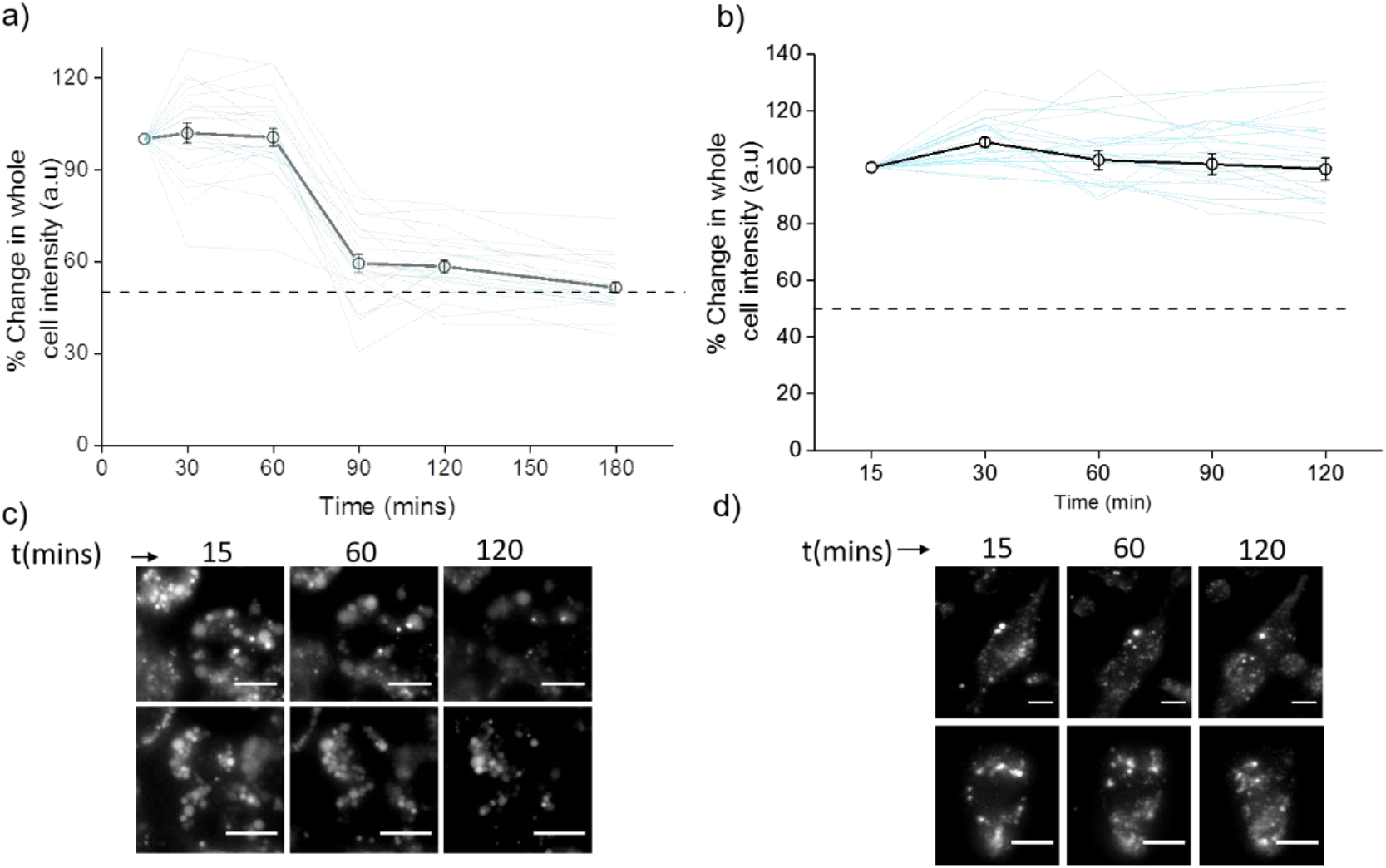
Plots showing mean whole cell intensity (black line) of *Clensor*A647 in cells as a function of indicated times in (a) J774A.1 cells and (b) THP-1 cells. Blue lines show traces of individual cells as a function of time. Error bars indicate s.e.m. (n ≥ 25 cells). Representative images of (a) J774A.1 cells and (b) THP-1 cells at indicated time points. Scale bar: 10 μm

To examine the stability of *Clensor in cellulo*, we pulse 500 nM *Clensor* for 30 min and chase for other 60 min to J774A.1 cells and THP-1 cells. The whole cell intensity of *Clensor_A647_* was unchanged upon 60 min chasing and decrease significantly from 60 min to 90 min for J774A.1 cell while no significant change was observed for THP-1 cells. The result indicates that the DNA device is stable for lysosomal pH measurement using our protocol (30 min pulse follow by 60 min chasing).

**Figure 4 figure supplementary 3:**
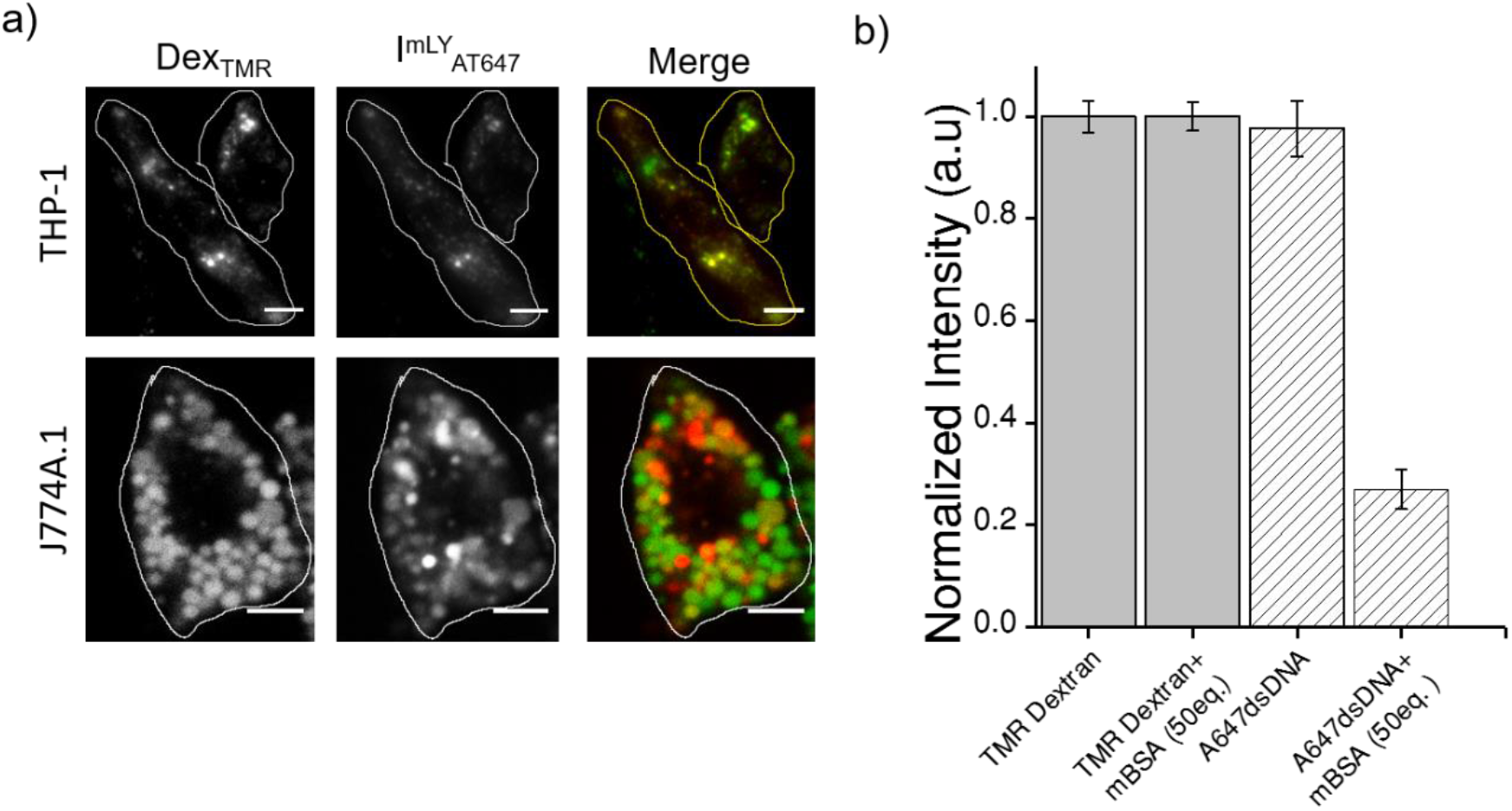
(a) Representative images showing colocalization of I^mLy^_AT647_ with TMR-Dex in J774A.1 and THP-1 macrophages (b) Macrophage labeling efficiency with *Clensor_A647_* (A647) in the absence or presence of 150 or 300 equivalents excess of maleylated bovine serum albumin (mBSA) in comparison to TMR Dextran. Error bars indicate s.e.m. (n = 20 cells).

To verify whether I^mLy^ was endocytosized to lysosome by anionic ligand binding receptor (ALBR) pathyway, cells were first pre-labelled with TMR-Dextran a marker of lysosome through fluid phase (0.5 mg/mL; 1 h pulse followed by 16 h chase). Cells were then pulse I^mLy^ for 30 min and chased for 60 min. Co-localization of I^mLy^ with TMR-Dextran was observed. Next, competition experiments with 30 equivalents of maleylated BSA (mBSA) – a well-known competitor for the anionic ligand, were performed. In the presence of mBSA, only the intensity of I^mLy^ was decreased significantly thus confirming that I^mLy^ was endocytosized *via* ALBR pathway.

**Figure 4 figure supplementary 4:**
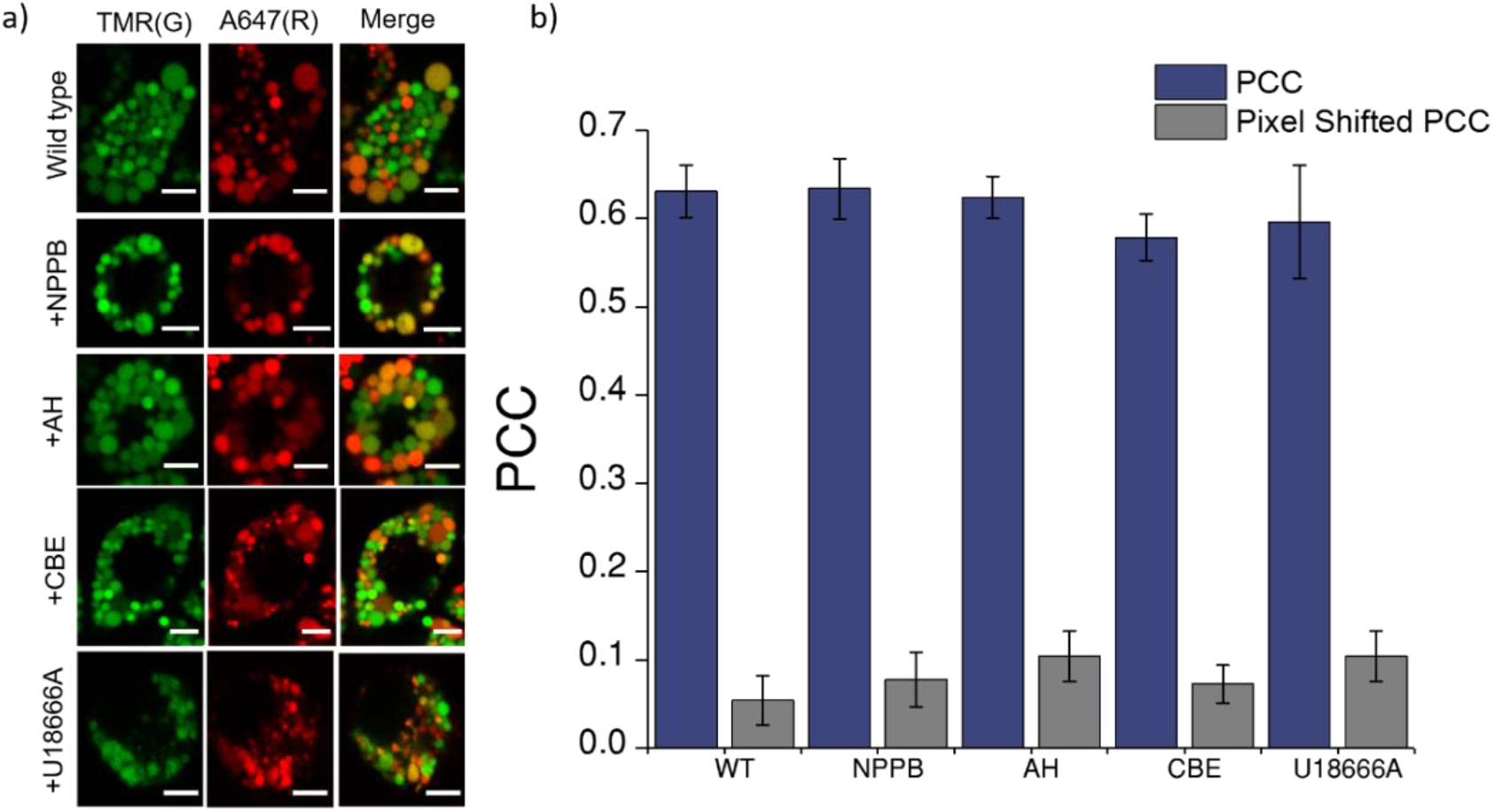
Co-localization of *Clensor* (red) with lysosomes labelled with TMR–dextran (green) in *J774A.1* cells treated with the indicated lysosomal enzyme inhibitors. (A) Representative images of lysosomes of *J774A.1* cells labelled with TMR dextran (TMR; G) and *Clensor* (Alexa 647; R) in the absence of any inhibitor, 50 μM NPPB, 10 μM Amitryptiline hydrochloride (AH) 400 μM of Conduritol b-epoxide, 10 μM of U18666A or 1 mM N⅛Cl. Scale bar, 5 μm c) Pearson’s correlation coefficient for the colocalization between *Clensor* and TMR Dextran.

To eliminate the possibility that trafficking defect in LSD may contributes to the observed reduction in lysosomal [Cl^−^]. We pre-labelled lysosomes with TMR-Dextran (0.5 mg/mL) using literature protocols, treated cells with U18666A and Conduritol β-epoxide (CBE) to induce Niemann Pick C and Gaucher’s cellular phenotype, only then pulse *Clensor* for 30 min, chased for 60 min, and scored for co-localization with TMR-Dextran. Amitriptyline (AH) and NPPB were added 30 min after the pulsing and chasing of *Clensor* to the lysosome. We observed that *Clensor* colocalized with lysosomes under each of these conditions, indicating that they do not suffer trafficking defects in these cell culture models of lysosomal storage diseases.

**Figure 4 figure supplementary 5:**
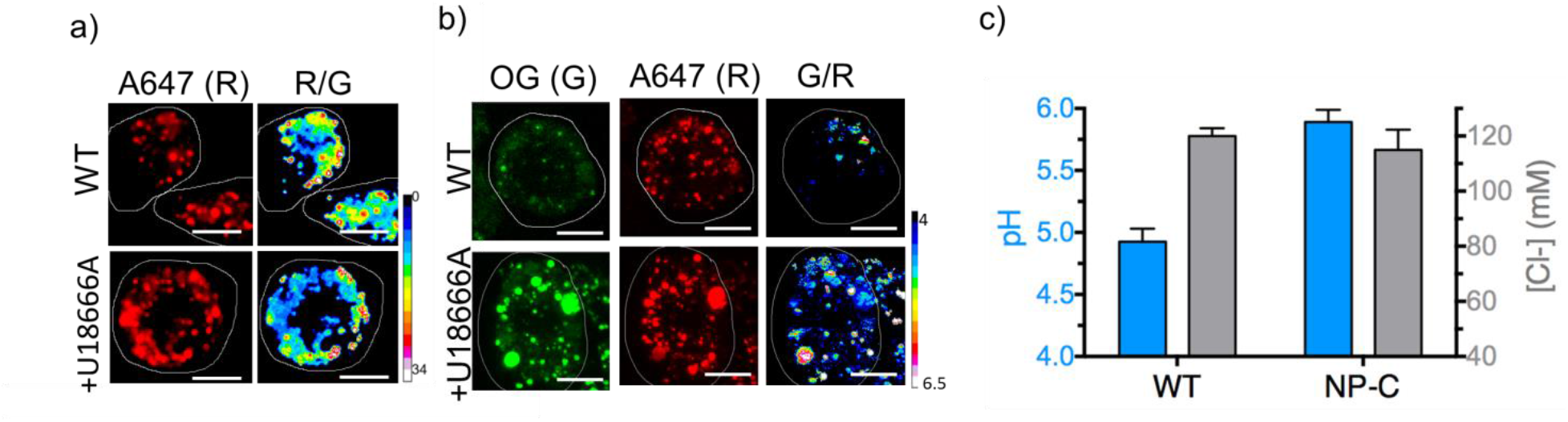
Representative pH and [Cl^−^] maps of *Clensor* in lysosomes of J774A.1 cells in the absence and presence of U18666A that gives a cell culture model of NP-C. (A) Representative images of lysosomes labeled with Clensor in the Alexa 647 (R) channel and pseudocolored R/G images are shown. (B) Representative images of lysosomes labeled with pH sensor I^mLy^ in the Oregon Green channel, red channel (R) and pseudocolored G/R images are shown. (C) pH and chloride levels in lysosomes in the presence and absence of U18666A. Error bars indicate s.e.m. (n = 10 cells, ≥50 endosomes). Scale bar: 10 μm

To directly compare the results from ncr-1 knockdowns that yielded a worm model of Niemann Pick C in Fig. 3, we also employed U18666A a selective inhibitor for NPC1 protein to induce NP-C cell model. We first confirmed that *Clensor* could traffic to the lysosome in this cell culture model (Fig. 4 figure supplementary 5). We found a dramatic lysosomal hypoacidification and no significant change of lysosomal [Cl^−^]. This matches our observation in a *C. elegans* model for NP-C.

**Figure 5 figure supplementary 1:**
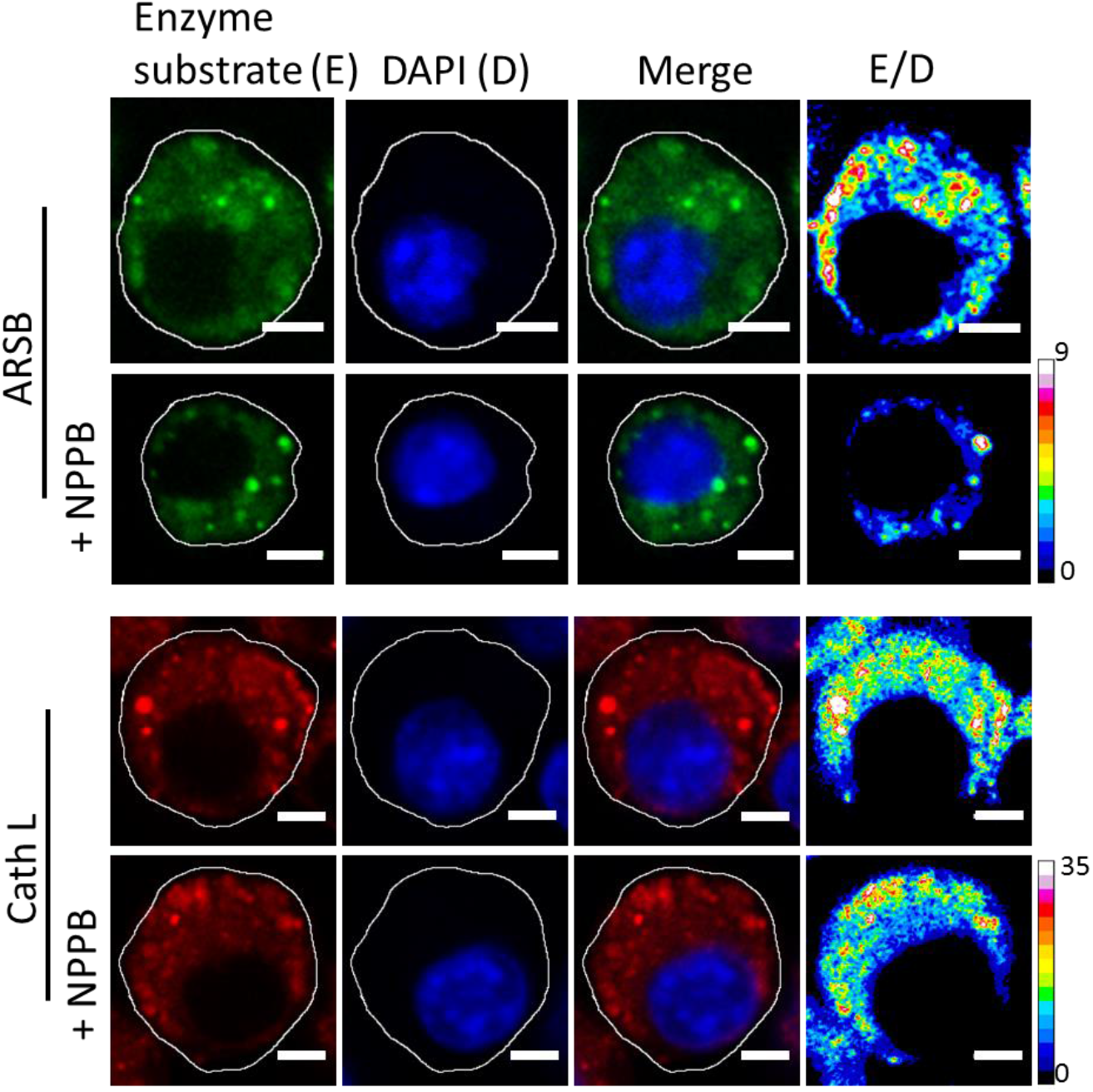
Fluorescence of cleaved substrates of ARSB and cathepsin L (E), DAPI (D) merge of E and D channels and respective pseudocolour E/D maps of J774A.1 cells with and without 50 μM NPPB. Scale bar: 5 μm.

To demonstrate chloride ion levels directly influence lysosome function, 50 μM of chloride channel inhibitor – NPPB was employed to decrease lysosomal chloride without affecting lysosomal pH (Fig. S13). Upon incubation of 50 μM of NPPB for 30 min, the activity of arylsulfatase B (ARSB) was reduced *ca*. 50% while the activity of its upstream enzyme cathepsin L which activates cathepsin C,(Dahl et al., 2001) is still unchanged. It reveals that decrease of chloride ions level directly attenuate cathepsin C activity.

**Figure 5 figure supplementary 2:**
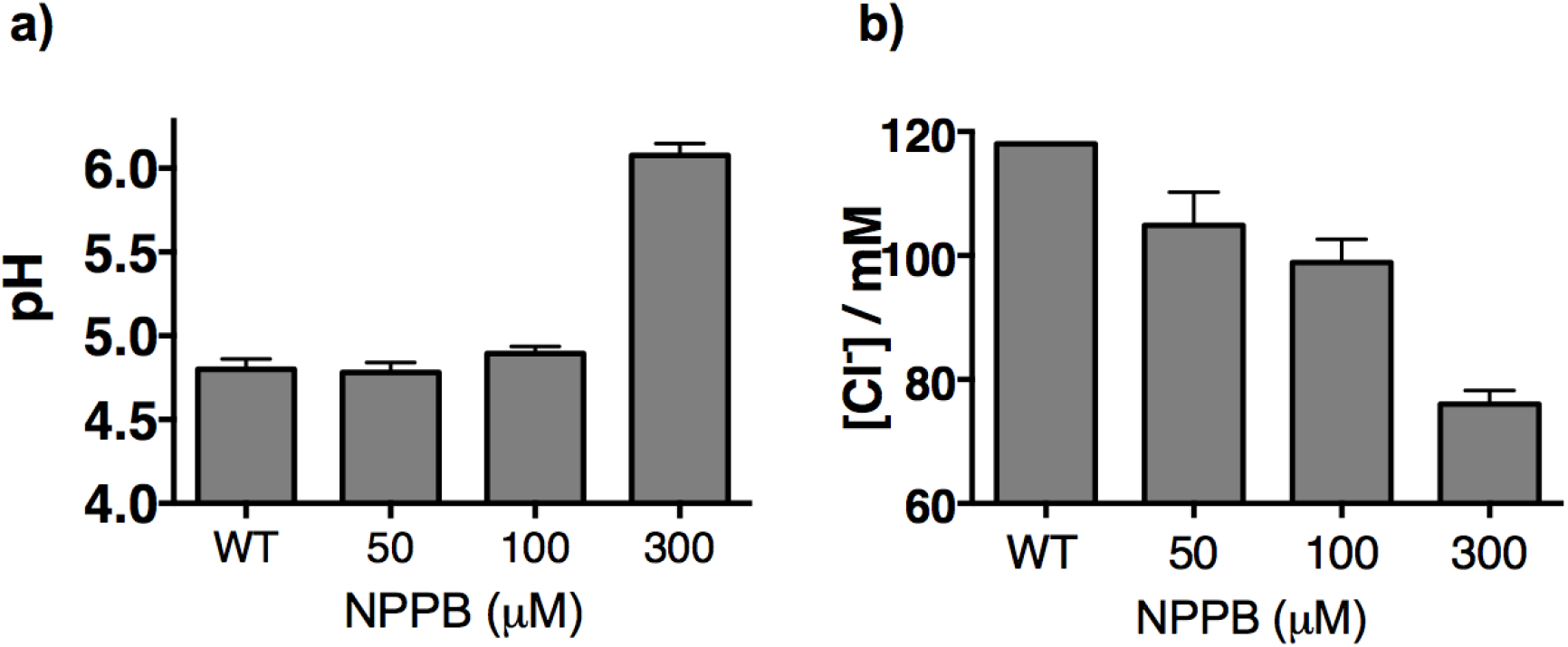
Lysosomal pH (a) and chloride (b) levels measured by I^mLy^ and *Clensor* in J774A.1 cells with increasing concentrations of NPPB.

Here we show that by varying doses of NPPB that at <100 μM of NPPB, we can selectively reduce lysosomal Cl-without changing in lysosomal pH. Consequently all further studies that use NPPB do so at 50 μM.

